# Exosomes promote axon outgrowth by engaging the Wnt-Planar Cell Polarity pathway

**DOI:** 10.1101/2023.10.28.564542

**Authors:** Samar Ahmad, Melanie Pye, Masahiro Narimatsu, Siyuan Song, Tania Christova, Jeffrey L. Wrana, Liliana Attisano

## Abstract

In neurons, the acquisition of a polarized morphology is achieved upon the outgrowth of a single axon from one of several neurites. Exosomes or small extracellular vesicles (sEVs) from diverse sources are known to promote the neurite outgrowth and thus may have therapeutic potential. However, the effect of fibroblast-derived exosomes on axon elongation in neurons of the central nervous system under growth permissive conditions remains unclear. Here, we show that fibroblast-derived sEVs promote axon outgrowth and a polarized neuronal morphology in mouse primary embryonic cortical neurons. Mechanistically, we demonstrate that the sEV-induced increase in axon outgrowth requires endogenous Wnts and core PCP components including Prickle, Vangl, Frizzled and Dishevelled. We demonstrate that sEVs are internalized by neurons, colocalize with Wnt7b and induce relocalization of Vangl2 to the distal axon during axon outgrowth. In contrast, sEVs derived from neurons or astrocytes do not promote axon outgrowth, while sEVs from activated astrocytes inhibit elongation. Thus, our data reveals that fibroblast-derived sEVs promote axon elongation through the Wnt-PCP pathway in a manner that is dependent on endogenous Wnts.

## Introduction

Neurons are the fundamental unit of the central nervous system (CNS) and exhibit a polarized morphology comprised of a single long axon and one or multiple dendrites. Sensory signals are received by dendrites and propagated down the length of the axon to neighboring neurons (Arimura & Kaibuchi, 2007; Namba et al., 2015; Schelski & Bradke, 2017; Takano et al., 2019). Thus, proper neuronal morphology is critical for synaptic transmission and cognitive functions of the brain. To acquire a polarized morphology, one neurite outgrows the other minor neurites and forms an axon, a process referred as neuronal polarization (Arimura & Kaibuchi, 2007; Namba et al., 2015; Schelski & Bradke, 2017; Takano et al., 2019). Axons differ in their growth potential such that during development axons grow rapidly but in the adult CNS, axons have lost the ability to grow (Bradke, 2022). Hence, the study of the mechanisms of axon outgrowth during development can provide insights into processes that need to be reactivated during axon regeneration in adults. Diverse extracellular factors have been reported to modulate neurite outgrowth and neuronal morphology (Arimura & Kaibuchi, 2007; Namba et al., 2015; Schelski & Bradke, 2017; Takano et al., 2019).

Exosomes are small extracellular vesicles (sEVs) that arise from multivesicular bodies (MVBs) within the endosomal pathway. Exosomes range in size from 30-150 nm and harbor a variety of biomolecules including proteins, DNA and RNAs (Colombo et al., 2014; Pegtel & Gould, 2019; van Niel et al., 2022). Exosomes regulate many processes including immune responses, wound healing and disease pathology (Zhang et al., 2019; Yates et al., 2022; Buzas, 2023). In the CNS, exosomes act as carriers for communication among neurons and glia, and promote neuroprotective functions (Chivet et al., 2013; Bahram Sangani et al., 2021; Gassama & Favereaux, 2021; Fan et al., 2022). The biological activities of exosomes depend upon the features of the cells from which they originate. For instance, exosomes from mesenchymal stem cells increase the number of neurites and promote neurite outgrowth in cortical neurons (Xin et al., 2012; Lopez-Verrilli et al., 2016; Zhang et al., 2017) while those from Schwann cells of the peripheral nervous system (PNS) promote axon regeneration in PNS neurons (Lopez-Verrilli et al., 2013). Peripheral nerve fibroblast-derived exosomes promote Schwann cell-mediated myelination of neurons (Zhao et al., 2022). Moreover, fibroblasts promote sorting of Schwann cells at an injury site that leads to axon regrowth (Parrinello et al., 2010). Thus, both fibroblasts and Schwann cells contribute to the remarkable regenerative ability of the PNS. In the context of growth inhibitory substrates, it has been shown that exosomes from fibroblasts can promote axon regeneration in cortical neurons (Tassew et al., 2017) suggesting that exosomes may have therapeutic potential. However, the direct effect of fibroblast-derived exosomes on axon outgrowth of CNS neurons during growth permissive conditions remains unclear. We previously showed that exosomes from fibroblasts can promote cell protrusion and motility in cancer cells (Luga et al., 2012). Given that core mechanisms of cytoskeletal remodeling are conserved during non-neuronal cell migration and neurite outgrowth in neurons, we sought to examine whether exosomes might similarly play a role in modulating neurite outgrowth. Thus, the current study is aimed at investigating the axon outgrowth promoting properties of fibroblast-derived exosomes.

Exosomes have previously been shown to mobilize autocrine Wnt-Planar cell polarity (PCP) signaling in breast cancer cells (Luga et al., 2012). PCP signaling was first described for its function in modulating tissue polarity by aligning cells in a single plane, perpendicular to the apical-basal axis (Goodrich & Strutt, 2011; Yang & Mlodzik, 2015; Devenport, 2016; Koca et al., 2022). In vertebrates, PCP signaling also functions in the development of inner ear, orientation of hair follicles and convergent extension (CE) movements during gastrulation and neural tube closure (Jones & Chen, 2007; Wallingford, 2012; Butler & Wallingford, 2017; Davey & Moens, 2017; Wang et al., 2019; Montcouquiol & Kelley, 2020). In the CNS, PCP modulates numerous developmental processes including differentiation of neural progenitor cells, neuronal migration, axon guidance and dendritic arborization (Goodrich, 2008; Davey & Moens, 2017; Hakanen et al., 2019; Zou, 2020). Core components of the PCP pathway are the seven-pass transmembrane Frizzled (Fzd) receptors, four-pass transmembrane Van-Gogh-like (Vangl) proteins, and cytoplasmic proteins including Prickle (Pk) and Dishevelled (Dvl) (Goodrich & Strutt, 2011; Yang & Mlodzik, 2015; Devenport, 2016; Koca et al., 2022). Unlike in Drosophila, Wnt morphogens play an important role in regulating PCP signaling in vertebrates, and may act as an instructive cue (Butler & Wallingford, 2017).

Wnts are lipid-modified secreted glycoproteins that act as morphogens to regulate processes during embryonic development including cell proliferation and fate, polarity and migration (Willert & Nusse, 2012; Nusse & Clevers, 2017; Steinhart & Angers, 2018; Mittermeier & Virshup, 2022; Rim et al., 2022). In neurons, Wnts are known to modulate dendritogenesis, axon guidance and remodelling, synaptogenesis, synaptic plasticity and differentiation (Salinas, 2012; Bonansco et al., 2022). In mammals, there are 19 Wnts that bind to different classes of cell surface receptors and trigger diverse cellular responses (Rim et al., 2022). Wnt receptors and co-receptors include members of Fzd, low-density lipoprotein related receptor (LRP) and receptor tyrosine kinases (RTKs) such as ROR, RYK, and PTK7 (MacDonald & He, 2012; Niehrs, 2012). Despite increasing knowledge about Wnt signaling, little is known about the role of Wnt-PCP pathway in axon outgrowth.

The current study was aimed at investigating the contribution of Wnt-PCP signaling in exosome-induced axon outgrowth. We show that sEVs derived from diverse fibroblast cell lines promote axon outgrowth that leads to acquisition of a polarized neuronal morphology. Mechanistically, we show that the sEV-induced increase in axon outgrowth requires neuronally-expressed Wnts and PCP components including Pk, Vangl, Fzd and Dvl. Importantly, we show that sEVs are internalized by neurons where they can colocalize with Wnt7b and also induce a shift in Vangl2 localization towards distal part of axon. In contrast to fibroblast-derived sEVs, those isolated from neurons or astrocytes do not alter axon outgrowth, while sEVs isolated from activated astrocytes inhibit axon outgrowth. Overall, the current study uncovers the potential of fibroblast-derived sEVs in activating axon outgrowth systems through PCP signaling, and may also indicate the therapeutic potential of sEVs in treating neuronal injury.

## Results

### sEVs promote acquisition of a polarized neuronal morphology by enhancing axon outgrowth

Dissociated primary mouse embryonic cortical neurons when freshly plated *in vitro* (Fig. S1A) give rise to short neurites, one of which grows rapidly to form an axon by 24-48 hours to generate a polarized neuron (Dotti et al., 1988; Arimura & Kaibuchi, 2007). Recent studies have demonstrated that exosomes isolated from mesenchymal stem cells can promote neurite outgrowth (Lopez-Verrilli et al., 2016; Zhang et al., 2017). Thus, we sought to explore the role of fibroblast derived EVs in neurite outgrowth. Following the guidelines of International Society for Extracellular vesicles (ISEV) (Welsh et al., 2024), the isolated vesicles are hereafter referred as sEVs, instead of exosomes. We first isolated and characterized sEVs from diverse mouse and human fibroblast cell culture supernatants including mouse L cells, mouse embryonic fibroblasts (MEFs), human dermal fibroblasts (HDFn), normal human lung fibroblasts (NHLF) and BJ foreskin fibroblasts using differential centrifugation (Fig. 1A). Analysis of the final pellet by immunoblotting revealed the presence of EV markers CD81, TSG101 and Flotillin1, and the absence of the endoplasmic reticulum (ER)-resident protein, Calnexin (CNX) (Fig. 1B). Particle size distribution analysis of the final pellet using nanoparticle tracking analysis (NTA) showed that the majority of particles were within a size range of 30-150 nm (Fig. 1C) that is characteristic of exosomes or sEVs (Colombo et al., 2014). Ultrastructural analysis by transmission electron microscopy (TEM) further showed intact round vesicles ranging in sizes of 30-150 nm in diameter, thus, confirming the identity and structural integrity of the isolated sEVs (Fig. 1D).

**Figure 1.**
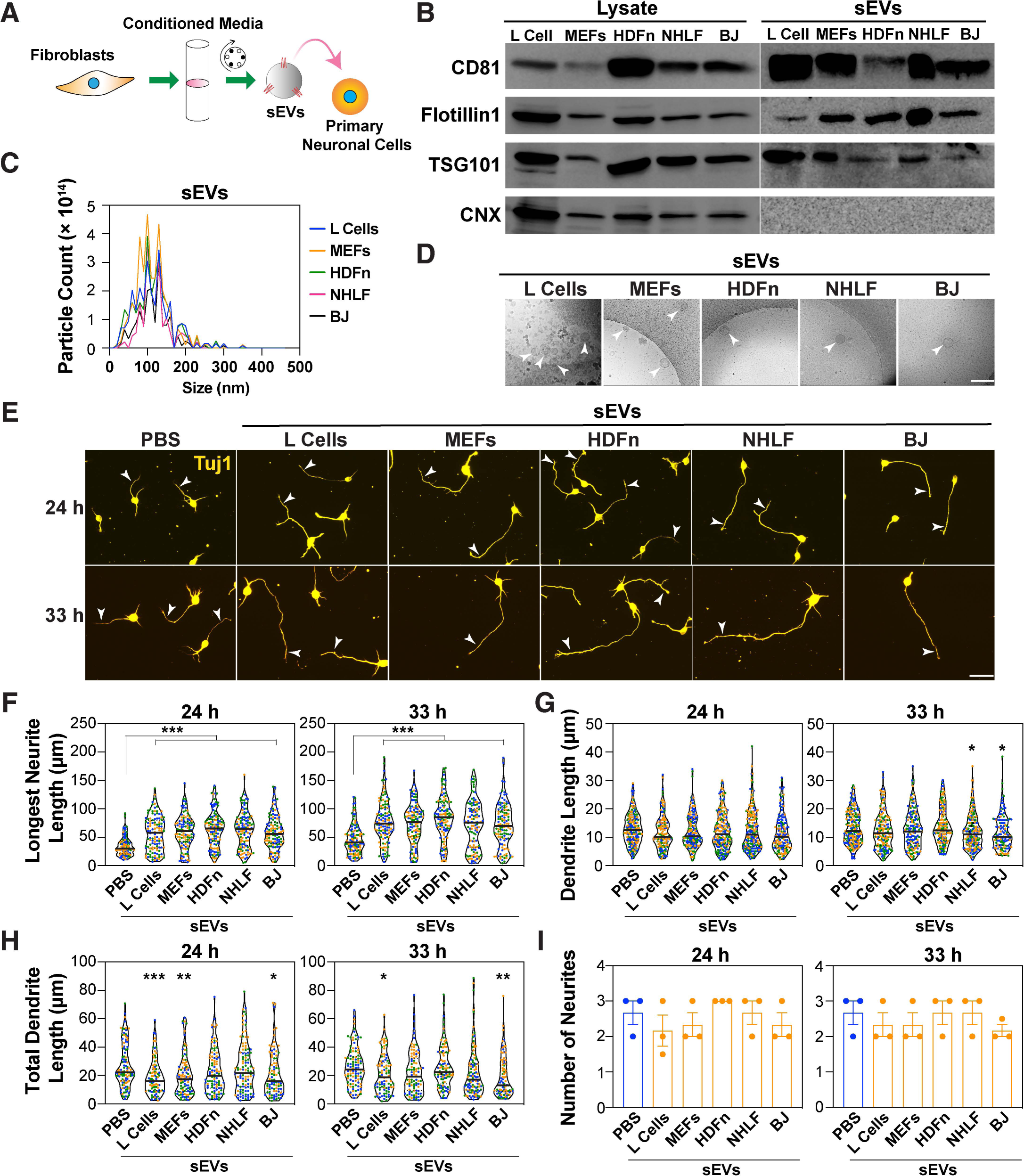
sEVs promote the growth of the prospective axon. **(A)** A schematic of the experimental set up. Mouse cortical neurons (E15.5-16.5) are treated with sEVs isolated from fibroblast-conditioned media (CM) using differential centrifugation. (**B)** Representative immunoblotting of cell lysates and sEV pellets (100,000 x g) from the indicated fibroblast cell lines for EV markers CD81, Flotillin1 and TSG101 and the ER protein, calnexin (CNX). (**C)** Nanoparticle tracking analysis (NTA) of differential centrifugation pellets. A representative plot indicating the particle size distribution from 3 independent purifications is shown. (**D)** Representative transmission electron microscopy (TEM) images of sEV-containing pellets. Arrowheads indicate round vesicles. Scale bar, 200 nm. (**E-I)** Cortical neurons were treated with sEVs (5 μg/mL) purified from the indicated fibroblast cell lines, 4 h after plating. Neurons were fixed at 24 and 33 h, and neuronal morphology was examined after staining for Tuj1. **(E)** Representative images are shown. Arrowheads mark the longest neurite. Scale bar, 40 μm. **(F-I)** The length of the longest neurite (prospective axon; **F**), individual neurite/dendrite lengths **(G)**, total dendrite length (longest neurite excluded; **H**) and total number of neurites **(I)** were quantified from a minimum of 90 neurons per condition from 3 independent experiments. Neurite lengths are plotted as a violin plot with values from each experiment distinctly colored and the median marked by a black line **(F, G, H)**. The number of neurites are plotted as the average of the median ± SEM **(I)**, where each dot represents the median from 30 neurons from one of the three independent experiments. Statistical significance: *p<0.05, **p<0.01, ***p<0.001 using one-way ANOVA with Dunnett’s post-test.

We next investigated the role of the fibroblast-derived sEVs on the neuronal morphology using dissociated E15.5 – 16.5 primary mouse embryonic cortical neurons. Neurons were treated 4 h after plating with sEVs for either 20 or 29 hours and were fixed and stained with Tuj1 antibody, which detects the neuron specific β-III tubulin. Quantification of neurite lengths revealed that sEVs from all five fibroblast lines enhanced the length of the longest neurite (ie the prospective axon) from a median length of ∼30 μm (in controls) to ∼60 μm at 24 h and from ∼40 μm (in controls) to ∼75 μm at 33 hours (Fig. 1E and 1F). In contrast, the individual lengths (∼10 μm) and the total combined length (∼20 μm) of the remaining neurites, excluding the longest, referred to as minor neurites and correspond to prospective dendrites, remained either unchanged or slightly reduced (Fig. 1G and 1H). The total number of neurites was also unaltered by sEV treatment (Fig. 1I). Similarly, when neurons were treated with varying concentrations of sEVs (0.05-10 μg/mL) obtained from all five fibroblast cell lines, a dose-dependent increase in the length of the longest neurite at both 24 and 33 hours was observed with the maximal effect occurring at 5 μg/mL (Fig. S1B-K). To verify that sEVs preferentially act on axons, neurons treated with sEVs from L cells were co-stained with the axonal marker, Tau-1 and the dendritic marker, MAP2. Quantification of Tau-1 positive axons confirmed that sEVs increased the length of axons (Fig. S2A and S2B), yielding results similar to those obtained when the length of the longest neurite, independent of Tau-1 staining was quantified in parallel (Fig. S2C). Similar to the data observed in Fig. 1G, the length of individual neurites (∼10 μm) positive for the dendritic marker MAP2 was either unaffected or slightly reduced by sEVs (Fig. S2D). Moreover, sEV treatment concordantly resulted in a significant increase in the percent of neurons with Tau-positive axons, a characteristic of a polarized neuron that was clearly evident within 24 h of plating (Fig. S2E). Importantly, the increase in the percent of Tau-positive axons could be observed as early as 12 hours after plating, indicating the rapid effect of sEVs (Fig. S2F and S2G). Taken together, these results demonstrate that sEVs derived from a variety of fibroblast lines, promote axon outgrowth and accordingly enhance the acquisition of a polarized phenotype.

The sEV-induced increase in axon outgrowth was lost when sEVs were incubated with detergent, which causes lysis of sEVs (Cvjetkovic et al., 2016; Bonsergent et al., 2021) or upon treatment with Proteinase K which digests surface accessible proteins (Fig. S3A and S3B). This indicates that while the axon-promoting activity is not found inside the sEVs, intact sEVs are required for function, suggesting that the active proteins might be expressed on the surface of sEVs. Next, to rule out a potential role for contaminating cell membranes or protein aggregates in mediating the effects on neuronal morphology, we undertook two different and more stringent exosome isolation protocols, where the final ultracentrifugation (100,000 x g) pellet of sEVs obtained from L cell supernatants was further subjected to either density gradient purification (Fig. S3) or size exclusion chromatography (Fig. S4). For density gradient separation, the resuspended pellet was applied to a discontinuous iodixanol gradient and then subjected to an additional ultracentrifugation (Fig. S3C). Characterization of the floating fractions by immunoblotting demonstrated that Fractions 6 – 8 were positive for the EV markers, CD81 and Flotillin1 (Fig. S3D) with the intensity of both markers peaking in Fraction 7 (Fig. S3E). Of note, Fraction 9 had the highest amount of total protein but very low levels of EV markers (Fig. S3E) indicating a successful enrichment of EV particles in Fractions 6 - 8. Particle size distribution analysis showed that the number of particles peaked at approximately 100 nm, which is a characteristic of exosomes or sEVs, and that Fraction 7 had the highest number of particles (Fig. S3F). Moreover, Fraction 7 (F7) had the highest particle/protein ratio (Fig. S3G), a parameter that is widely used to measure the purity of EVs (Webber & Clayton, 2013; Thery et al., 2018). Treatment of dissociated neurons with these purified sEVs from the F7 fraction revealed that the length of the longest neurite (ie the prospective axon) was increased to levels similar to that observed for the initial ultracentrifugation pellet (sEVs) at all time points (Fig. S3H and S3I).

Next, we used size exclusion chromatography as an alternative approach to purify sEVs from the ultracentrifugation (100,000 x g) pellets (Fig. S4A). Characterization of the isolated fractions revealed the presence of the EV markers, CD81, Flotillin1 and Tsg101 but not the ER protein, Calnexin (CNX) in fractions 7 – 12 (Fig. S4B), with particle size peaking at a diameter of approximately 100 nm with fraction F7 having the highest particle/protein ratio (Fig. S4C and D) which is characteristic of exosomes or sEVs. Treatment of dissociated neurons with F7 purified sEVs revealed that the length of the longest neurite (ie the prospective axon) was increased to levels similar to that observed for the initial ultracentrifugation pellet (sEVs) at both 24 and 33 hours (Fig. S4E and S4F) as observed previously (Fig. 1E, 1F and S1). Of note, outgrowth induced by sEVs isolated using either purification method was easily detected as early as 12 hours (ie 8 h after sEV addition), highlighting the rapid nature of the response (Fig. S3H, S3I, S4E and S4F).

Altogether, these observations suggest that the sEVs isolated by ultracentrifugation (100,000 x g pellet) are responsible for promoting axon outgrowth. Since the responses induced by these pelleted sEVs were similar to that achieved after more extensive purifications, we used this ultracentrifuged pellet as a source of sEVs for all subsequent experiments.

### The PCP pathway is required for sEV-induced growth of the prospective axon

Previous work has demonstrated that in cancer cells, motility induced by exogenously added fibroblast-derived exosomes is mediated by the Wnt planar cell polarity pathway (PCP) in recipient cells (Luga et al., 2012). Prickle (Pk) and Vangl, core PCP components, distribute asymmetrically to the proximal side in planar polarized epithelial cells (Yang & Mlodzik, 2015; Butler & Wallingford, 2017) and are also known to modulate neuronal morphology (Dreyer et al., 2022; Radaszkiewicz et al., 2023). This prompted us to investigate whether core PCP components might be required for sEV-induced axon outgrowth. To study this, we used a combination of siRNA-mediated knockdown and mouse genetic models to explore the role of these core PCP components in dissociated primary cortical neurons. There are two Prickle genes, Prickle 1 and Prickle 2 (Pk1 and Pk2), expressed in cortical neurons (Fig. S5A). To test for the requirement for Prickle, neurons were co-transfected with siRNAs targeting Pk1 and/or Pk2 along with plasmid, encoding GFP to visualize transfected cells. Analysis of GFP positive neurons in which expression of Pk1 and Pk2 alone or together was abrogated, revealed that neurons were comprised solely of minor neurites (mean length ∼10 μM), none of which were elongated upon treatment with sEVs (Fig. 2A and 2B). Knockdown efficiency was confirmed by qPCR after FACS sorting for GFP positive neurons (Fig. 2C). As an alternative to this transient approach, we explored the consequences on neuronal morphology using dissociated neurons isolated from mice with genetic loss of Pk1 or Pk2. While Pk2 knockout mice are viable (Tao et al., 2011), Pk1 knockout mice display early embryonic lethality (Tao et al., 2009). Thus, we generated mice with single floxed Pk alleles (Pk1^f/f^ and Pk2^f/f^) using targeted homologous recombination (Fig. S5B), which were then crossed with each other to obtain mice with double (Pk1^f/f^;Pk2^f/f^) floxed Pk alleles. Both single and double floxed mice, after removal of neomycin cassette, were viable, fertile, and exhibited no overt anatomical defects. Conditional knockout mice were then generated by crossing with Nestin-Cre mice to achieve loss of Pk1/2 in the developing brain. Loss of Pk1/2 expression in the cortex of Pk1^f/f^;Pk2^f/f^;Nestin-Cre E15.5 embryos was confirmed by immunofluorescence microscopy using Pk1 and Pk2 antibodies (Fig. S5C and S5D). Consistent with the siRNA results, analysis of cortical neurons (E15.5-16.5) isolated from mice with a Nestin-Cre conditional deletion of either Pk1 alone, Pk2 alone or both (Pk1^f/f^;Pk2^+/+^;Nestin-Cre, Pk1^+/+^;Pk2^f/f^;Nestin-Cre or Pk1^f/f^;Pk2^f/f^;Nestin-Cre, respectively) revealed the presence of only short neurites that failed to elongate in response to sEV treatment in contrast to the Pk1^+/+^;Pk2^f/f^ or Pk1^+/+^; Pk2^+/+^;Nestin-Cre controls (Fig. 2D and S5E).

**Figure 2.**
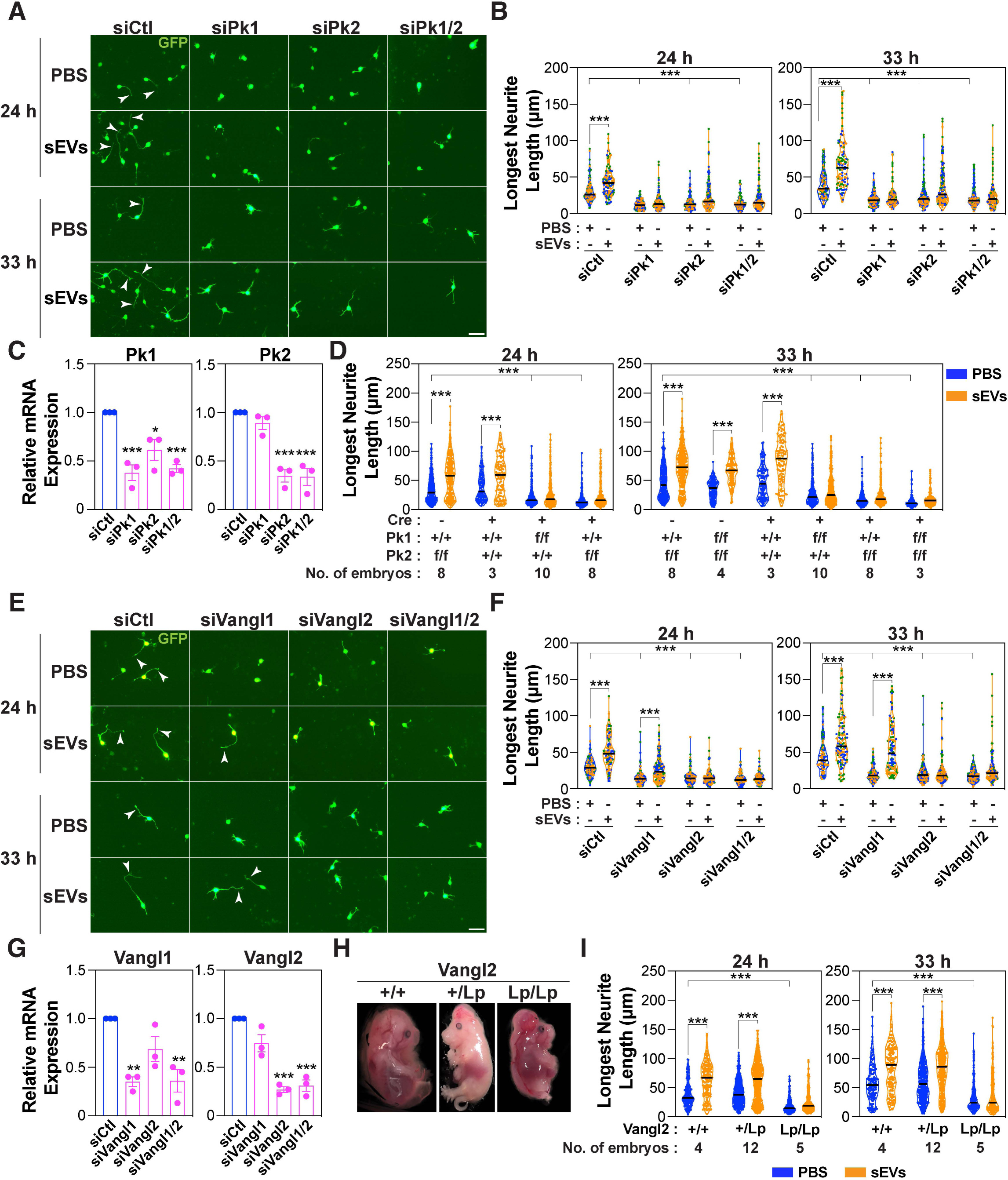
The PCP components, Pk1/2 and Vangl2, are required for sEV-induced growth of the longest neurite. **(A-C, E-G)** Pk1/2 and Vangl2 promote sEV-induced neurite outgrowth. Dissociated E15.5-16.5 mouse cortical neurons were electroporated with siRNA against Pk1 (siPk1) and Pk2 (siPk2) (**A-C**), or Vangl1 (siVangl1) and Vangl2 (siVangl2) (**E-G**) individually or in combination or with siControl (siCtl) along with a GFP-expressing plasmid and were treated with sEVs (5 μg/mL) from L cells, 4 h after plating. Neurons were fixed at 24 and 33 h, and neuronal morphology was examined in GFP-positive neurons. (**A, E**) Representative images are shown. Arrowheads mark the longest neurite. Scale bar, 40 μm. (**B, F**) The length of the longest neurites was quantified. (**C, G)** Knockdown efficiency for Pk1/Pk2 (**C**) and Vangl1/2 (**G**) was determined in GFP-positive neurons isolated by FACS. Relative mRNA expression was determined by qPCR. (**D, I)** Pk1/2 and Vangl2 promote sEV-induced neurite outgrowth in mutant mouse models. Cortical neurons (E15.5-16.5) were isolated from Pk1 and Pk2 conditional knockout mice obtained by crossing Pk floxed mice with a Nestin-Cre line **(D)** or Vangl2 mutant littermates obtained by crossing of heterozygous loop-tail mutants (Vangl2^+/Lp^) **(I).** Neurons were treated with sEVs from L cells, 4 h after plating, fixed at 24 and 33 h and morphology examined in Tuj1 stained neurons. The length of the longest neurite was quantified from 40 neurons per embryo, and total number of embryos analyzed are indicated below the genotypes. Violin plots with the median marked by a black line are shown. **(H)** Loop-tail embryos exhibit an open neural tube. Representative images are shown from a minimum 4 independent experiments. In siRNA experiments, neurite lengths are quantified from a minimum of 90 neurons per condition from 3 independent experiments and plotted as a violin plot with values from each experiment distinctly colored and the median marked by a black line **(B, F**). For qPCR plots, data is presented as the mean ± SEM from 3 independent experiments **(C, G)**. Statistical significance: *p<0.05, **p<0.01, ***p<0.001 using one-way ANOVA with Dunnett’s post-test **(C, G)**, or two-way ANOVA with Tukey’s post-test **(B, D, F, I)**.

We next explored the contribution of the core PCP tetraspanin proteins, Vangl1 and Vangl2, which regulate diverse aspects of neuronal development (Dreyer et al., 2022). Abrogation of Vangl2 expression using siRNAs strongly decreased sEV-mediated induction of outgrowth of the longest neurite (Fig. 2E-G). Interestingly, in the case of siVangl1, while the length of the longest neurite was reduced, sEVs maintained the ability to promote elongation (Fig. 2E-G), suggesting that Vangl2, but not Vangl1 is important for the sEV-mediated effects. Loop-tail mice harbor a S464N point mutation in the Vangl2 allele (Kibar et al., 2001; Murdoch et al., 2001) that results in reduced Vangl2 protein levels and defective translocation to plasma membrane (Torban et al., 2007; Merte et al., 2010; Belotti et al., 2012). Morphological examination showed that the heterozygous loop-tail embryos (Vangl2^+/Lp^) exhibited a curly tail with partial or no neural tube defects, whereas homozygous loop-tail embryos (Vangl2^Lp/Lp^) exhibited a completely open neural tube (Fig. 2H) as previously reported (Kibar et al., 2001). Consistent with the siRNA results, dissociated cortical neurons isolated from E15.5 – 16.5 Vangl2^Lp/Lp^ embryos, but not wild-type or heterozygotes (Lp/+) displayed short neurites (∼ 10 μM in length) and a loss of sEV-induced outgrowth of the longest neurite at both 24 and 33 hours (Fig. 2I and S5F).

Smurf1 and Smurf2 are E3 ubiquitin ligases that modulate PCP signaling to control convergent extension, cell protrusion and motility (Wang et al., 2003; Sahai et al., 2007; Narimatsu et al., 2009; Luga et al., 2012) and have also been shown to play a key role in regulating neuronal morphology (Vohra et al., 2007; Cheng et al., 2011; Kannan et al., 2012). Since cortical neurons express both Smurf1 and Smurf2 as determined by RNA sequencing analysis (Fig. S5A), we next explored whether Smurfs might have a role in sEV-induced axon outgrowth. For this, we examined the contribution of Smurf1 and/or Smurf2 using dissociated neurons isolated from Smurf knockout mice. Smurf1^+/−^;Smurf2^+/−^ double heterozygotes (Narimatsu et al., 2009) were crossbred to generate offspring with varying Smurf1/2 genotypes. Analysis of dissociated cortical neurons revealed that complete loss of either Smurf1 alone (Smurf1^−/−^; Smurf2^+/+^) or Smurf2 alone (Smurf1^+/+^;Smurf2^−/−^) yielded neurons that harboured only short neurites (∼ 10 μM in length) and that sEV-induced outgrowth of the longest neurite was prevented in contrast to wild-type or heterozygote (Smurf1^+/−^;Smurf2^+/−^) littermates (Fig. S5G and S5H). These results suggest that Smurf1 and Smurf2 act in a non-redundant manner or that a minimal level of combined Smurf1/2 activity is required for neurite outgrowth.

Both PCP and canonical β-catenin signaling is initiated upon binding of Wnt ligands to one of 10 members of the seven-pass transmembrane Frizzled family of receptors (Fzd) and is then propagated intracellularly through one of three Dishevelleds (Dvls) (van Amerongen et al., 2008; Gao & Chen, 2010; MacDonald & He, 2012; Yang & Mlodzik, 2015; Wang et al., 2016; Sharma et al., 2018). Thus, we next explored which Fzds and Dvls are required for sEV-mediated axon outgrowth. Cortical neurons treated with a pan-Fzd blocking antibody (F2.A) that recognizes multiple Fzds, including Fzd1, 2, 4, 5, 7 and 8 (Pavlovic et al., 2018; Tao et al., 2019) displayed only minor neurites that failed to elongate in response to sEVs (Fig. 3A and 3B). Treatment of cortical neurons with F2.A at a later time point had no effect on the axon length, however, the sEV-induced increase in the axon length was impaired, indicating that Fzds are required to mediate sEV effects (Fig. S6A and S6B). Mouse E15.5-16.5 cortical neurons express Fzd1, 2, 3, 7, 8 and 9 with Fzd1 and Fzd3 displaying the highest levels (Fig. S5A), thus we next explored the individual requirement for these receptors using siRNAs. Quantification revealed that individual loss of Fzd1, 3 or 9 yielded neurons with minor neurites that failed to elongate in response to sEVs while loss of Fzd7 only blocked sEV-mediated outgrowth (Fig. 3C, S6C and S6D). Loss of Fzd2 or Fzd8 had no effect, though equivalent knockdown efficiency was achieved (Fig. S6D). These results indicate that multiple but specific Fzds (Fzd 1, 3 and 9) are required for basal elongation, whereas only Fzd7 is specifically required for sEV-mediated outgrowth of the prospective axon. All three Dishevelleds (Dvl1, 2 and 3) are expressed in cortical neurons with Dvl1 being expressed at the highest level (Fig. S5A). Analysis of the requirement for Dvls using siRNAs revealed that only Dvl2 was required for sEV-induced outgrowth (Fig. 3D, S7A and S7B).

**Figure 3.**
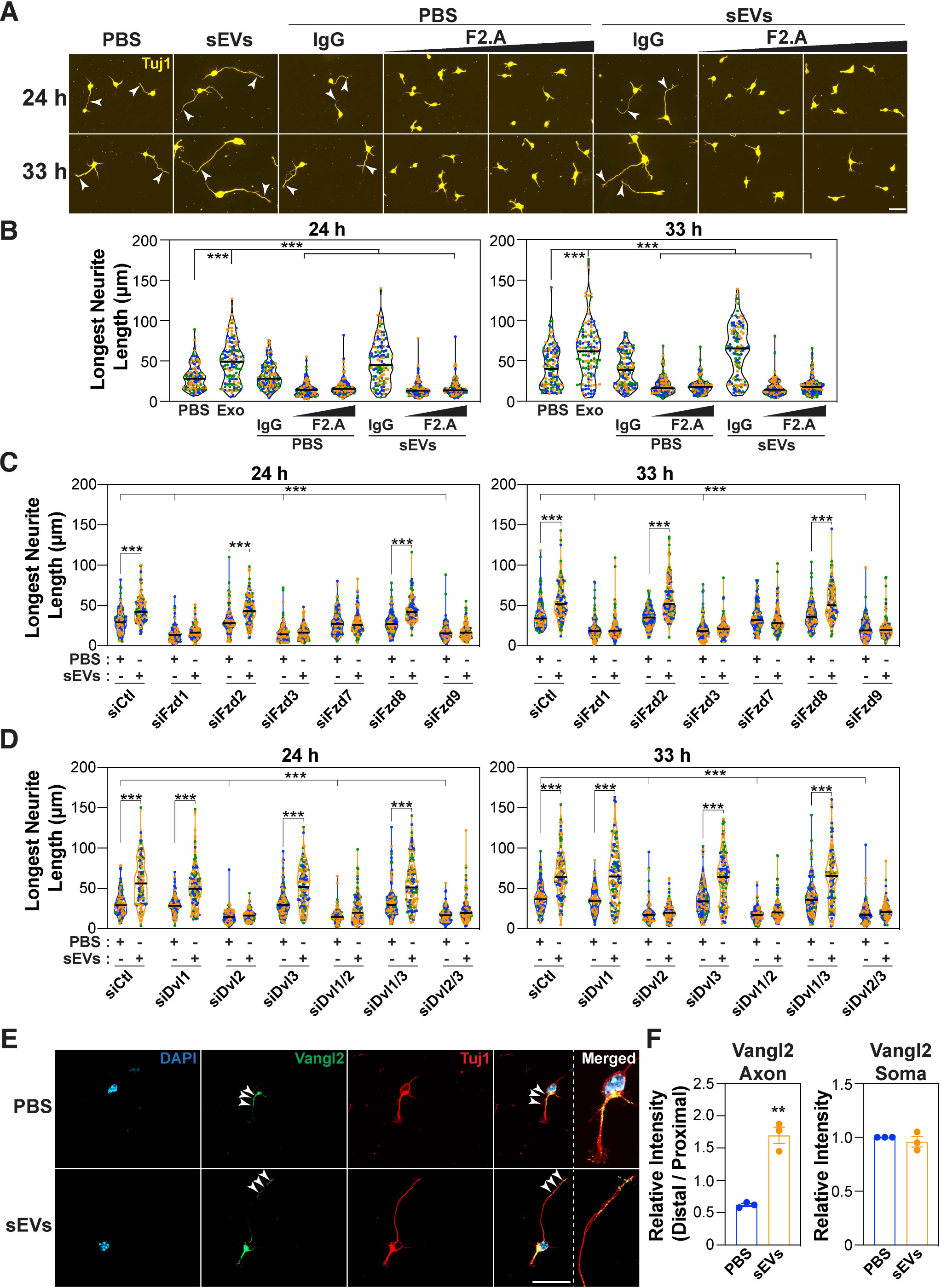
Fzds and Dvls are required for sEV-induced growth of the longest neurite. **(A-D)** Dissociated E15.5-16.5 mouse cortical neurons were treated with sEVs (5 μg/mL) from L cells, 4 h after plating. **(A-B)** Neurons were co-treated with IgG or a Fzd blocking antibody, F2.A (at 50 nM and 100 nM) along with sEVs. **(C-D)** Neurons were electroporated with siRNA against Fzds (siFzds) **(C)**, or Dvls (siDvl) **(D)** or siControl in combination with a GFP-expressing plasmid prior to the addition of sEVs. Neurons were fixed at 24 and 33 h, and neuronal morphology was examined in Tuj1 stained **(A, B)** or GFP-positive neurons **(C, D)** with representative images shown. Arrowheads mark the longest neurite. Scale bar, 40 μm. **(B-D)** The length of the longest neurite was quantified. **E)** sEVs promote localization of Vangl2 to the distal axon. Cortical neurons were treated with sEVs from L cells, 4 h after plating and fixed after 24 h. Representative confocal images of neurons stained with DAPI (blue), Vangl2 (green) and Tuj1 (red) are shown. Arrowheads mark the Vangl2 localization. (**F)** The ratio of distal/proximal Vangl2 intensity and the relative intensity of Vangl2 in soma is quantified from 30 neurons from 3 independent experiments. Neurite lengths are quantified from a minimum of 90 neurons per condition from 3 independent experiments and plotted as a violin plot with values from each experiment distinctly colored and the median marked by a black line **(B, C, D)**. For Vangl2 plots, data is presented as the mean ± SEM from 3 independent experiments **(F)**. Statistical significance: **p<0.01, ***p<0.001 using unpaired t-test **(F)**, one-way ANOVA with Dunnett’s post-test **(B)**, or two-way ANOVA with Tukey’s post-test **(C, D)**.

Individual Fzds and Dvls function in both PCP and β-catenin pathways in a complex and highly redundant manner thus it is challenging to associate specific Fzds/Dvls to a particular pathway (Gao & Chen, 2010; Mlodzik, 2016; Wang et al., 2016). However it is worth noting that evidence of Fzd7 and Dvl2, both of which we observed to be specifically required for sEV-induced growth of the prospective axon, have been reported to function in PCP pathways (Zou, 2020; Pascual-Vargas & Salinas, 2021). Taken together, these observations suggest that core PCP components are required for basal neurite elongation and that they are required for sEV-induced growth of the longest neurite.

### Vangl2 localizes asymmetrically during sEV-induced growth of the longest neurite

Prior work has revealed that Wnt5a-mediated activation of the PCP pathway in dissociated commissural neurons results in relocalization of Vangl2 to the growth cone (Shafer et al., 2011). Thus, we next explored whether sEVs might similarly promote relocalization of Vangl2. In control (PBS) or sEV-treated embryonic cortical neurons, Vangl2 was distributed in both the somal membrane and in the axon but in distinct patterns (Fig. 3E). Quantification of Vangl2 distribution within the axon shaft revealed that in controls, a preferential localization in the proximal half of the axon was observed (Fig. 3F), that was independent of axon length (Fig. S7C). Remarkably, in neurons treated with sEVs for 20 h, Vangl2 relocalized to the distal end of the axon (distal to proximal intensity ratio of ∼1.6 versus ∼0.6 in controls) and was often detected within the growth cone (Fig. 3F). No overt change in Vangl2 localization in the somal membrane was observed. This re-localization of Vangl2 to the distal axon and growth cone suggests that sEVs engage PCP components to promote axon growth.

### Neuronal Wnts mediate sEV induced growth of the longest neurite

Wnts, including Wnt5a, Wnt7a and Wnt7b can promote axon elongation, neurogenesis and axon guidance, respectively, via the PCP pathway (Salinas, 2012; Bonansco et al., 2022). As exosomes are known to carry Wnts (Routledge & Scholpp, 2019; Gross, 2021; Mittermeier & Virshup, 2022), we next explored whether the presence of Wnt ligands in isolated exosomes could contribute to the observed outgrowth of neurites. We first abrogated the expression of Porcupine, an acyltransferase essential for Wnt lipidation and secretion (Mikels & Nusse, 2006; Takada et al., 2006), in the exosome-producing L cells using siRNAs and isolated the sEVs (Fig. 4A). Abrogation of Porcupine expression had no effect on the overall number of sEVs secreted or on the levels of the EV markers, CD81, Flotillin1 and TSG101 as indicated by immunoblotting (Fig. S7D-G). Moreover, ultrastructural analysis by TEM revealed intact round, exosome-like vesicles in both the control and Porcupine knockdown samples (Fig. S7H). When applied to cortical neurons, sEVs isolated from Porcupine-deficient L cells were as active as control sEVs in promoting outgrowth of the longest neurite (Fig. 4B-D). Immunodetection of endogenous Wnt proteins in wild type L-cells is challenging, thus we confirmed that siPorcupine inhibits Wnt secretion using L-cells stably expressing Wnt5a (Fig. S7I-K). These results thereby indicate that the activity of L cell-derived sEVs was not due to the presence of Wnts within the sEVs.

**Figure 4.**
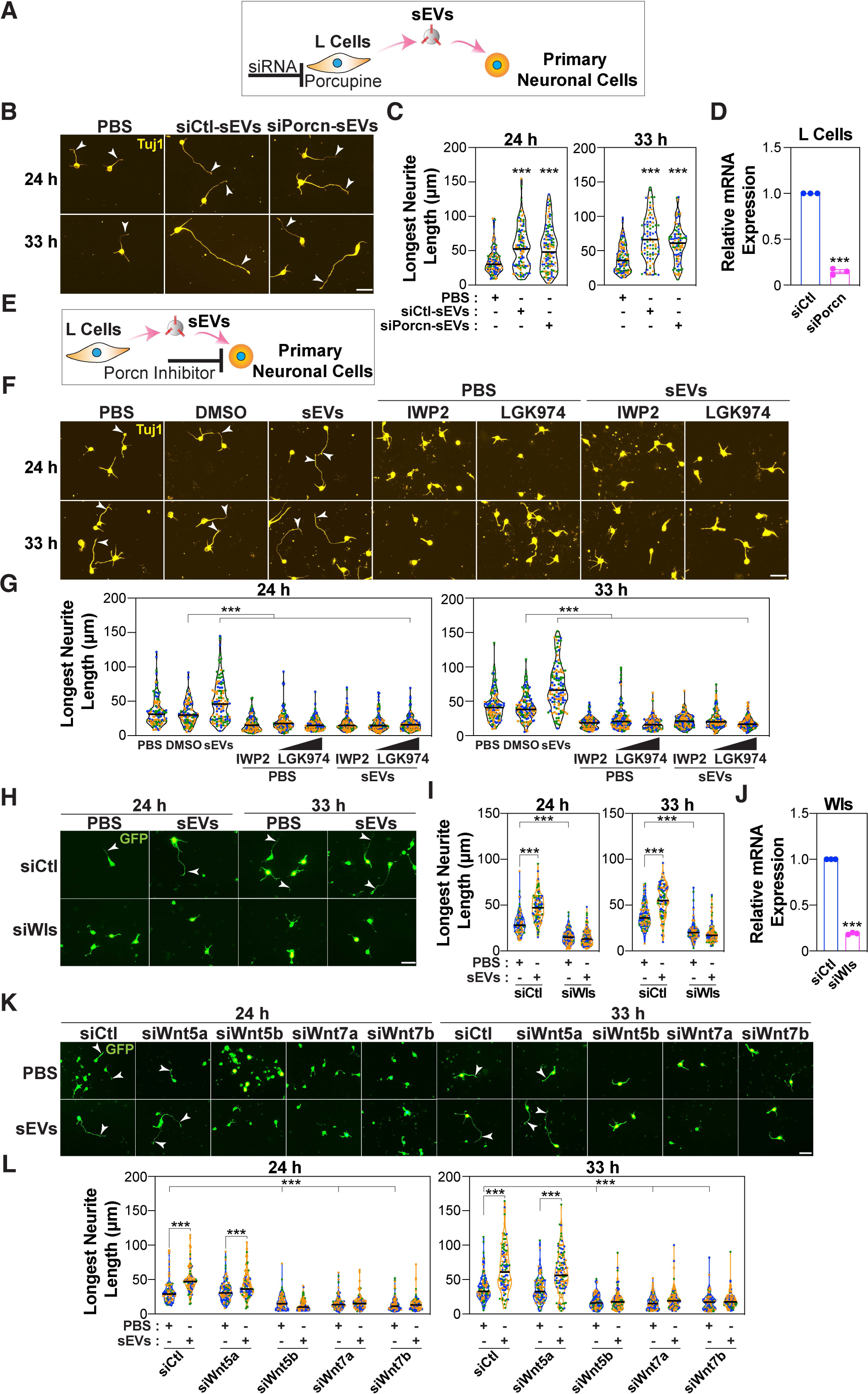
Neuronal Wnts mediates sEV-induced growth of the longest neurite. **(A, E)** Schematics, illustrating the experimental set up. **(A-G)** Dissociated E15.5-16.5 mouse cortical neurons are treated with sEVs isolated from L cells transfected with siCtl or siPorcupine **(A-D)** or with sEVs isolated from regular L cells. Neurons were treated with PBS as control and with porcupine (Porcn) inhibitors, IWP2 (10 μM) or LGK974 (1 and 5 μM) **(E-G)**, 4 h after plating and were co-incubated with sEVs. (**H-L)** Cortical neurons were electroporated with siRNAs against Wls (siWls) **(H, I)** or Wnts (siWnts) **(K, L)** or siControl (siCtl) along with a GFP-expressing plasmid and then treated with sEVs (5 μg/mL) from L cells, 4 h after plating. Neurons were fixed at 24 and 33 h, and neuronal morphology was examined in Tuj1 stained neurons **(B, C, F, G)** or GFP-positive neurons for siRNA experiments **(H, I, K, L)**. Representative images **(B, F, H, K)** and quantifications of the longest neurite **(C, G, I, L)** are shown. (**D, J)** Knockdown efficiency for porcupine **(D)** and Wls **(J)** was determined in L cells and GFP-positive neurons isolated by FACS, respectively. Relative mRNA expression was determined by qPCR. Neurite lengths are quantified from a minimum of 90 neurons per condition from 3 independent experiments and plotted as a violin plot with values from each experiment distinctly colored and the median marked by a black line **(C, G, I, L)**. For all other plots, data is presented as the mean ± SEM from 3 independent experiments **(D, J)**. Statistical significance: ***p<0.001 using unpaired t-test **(D, J)**, one-way ANOVA with Dunnett’s post-test **(C, G)**, or two-way ANOVA with Tukey’s post-test **(I, L)**.

We next sought to determine whether L cell-secreted exosomes might modulate autocrine Wnt signaling, a process whereby secreted Wnts bind to cell-surface receptors and thereby activate Wnt signaling in neurons to promote neurite outgrowth. To investigate this, we blocked the ability of neurons to secrete Wnts, thus attenuating the autocrine signaling, by chemically inhibiting the activity of Porcupine using IWP2 or LGK974 (Chen et al., 2009; Liu et al., 2013) (Fig. 4E). As endogenous Wnt proteins are difficult to detected in neurons, we confirmed that IWP2 blocked Wnt secretion using L-Wnt5a cells (Fig. S7L) as described above for siPorcupine (Fig. S7-I-K). We observed that treatment of cortical neurons with Porcupine inhibitors 4 h after plating, prevented sEV-mediated neurite outgrowth (Fig. 4F and G). In contrast, treatment with IWP2 at 24 h after plating, had no effect on basal axon lengths but impaired the sEV-induced axon outgrowth, thus indicating that neuronal Wnts are required to mediate the effect of sEVs (Fig. S8A and S8B). The Wnt carrier protein, Wntless (Wls) is required for Wnt secretion (Banziger et al., 2006), and abrogating expression of Wls using siRNAs, similarly prevented sEV-induced neurite outgrowth (Fig. 4H-J). Thus, blocking pathways essential for Wnt secretion prevents sEV-induced growth of the longest neurite, indicating that neuronally-expressed endogenous Wnts are necessary for sEV function.

Analysis of RNA sequencing results revealed that cortical neurons express Wnt5a, Wnt5b, Wnt7a and Wnt7b (Fig. S5A). We verified by qPCR that these Wnts were expressed in neurons and that expression levels remain constant throughout the time frame of the experiment (Fig. S8C). Thus, to determine which Wnts are important for neurite outgrowth, the expression of each Wnt was individually abrogated using siRNAs (Fig. S8D). Neurons lacking Wnt5b, Wnt7a or Wnt7b but not Wnt5a were comprised solely of minor neurites, none of which elongated upon treatment with sEVs (Fig. 4K and 4L). Altogether, these observations suggest that exogenously added sEVs might engage endogenous Wnts (Wnt5b, Wnt7a and Wnt7b) to promote growth of the longest neurite.

### sEVs associate with Wnts during neurite outgrowth

To explore sEV uptake in neurons, we used a compartmentalized microfluidic chamber device (Taylor et al., 2005) comprised of a set of proximal compartments, where neurons are seeded, and a set of distal compartments where axons emerge. The compartments are connected by a microgroove barrier that enables a gradient based fluidic separation due to the hydrostatic pressure (Fig. 5A). Plated neurons were grown for 5 days to allow axons to reach the distal compartment. sEVs were added to either compartment for 24 hours and then neurons were fixed and stained for Tuj1. Addition of sEVs to the proximal compartment containing the cell bodies but not to the distal compartment containing axon tips, promoted elongation of neurites (Fig. 5B and 5C).

**Figure 5.**
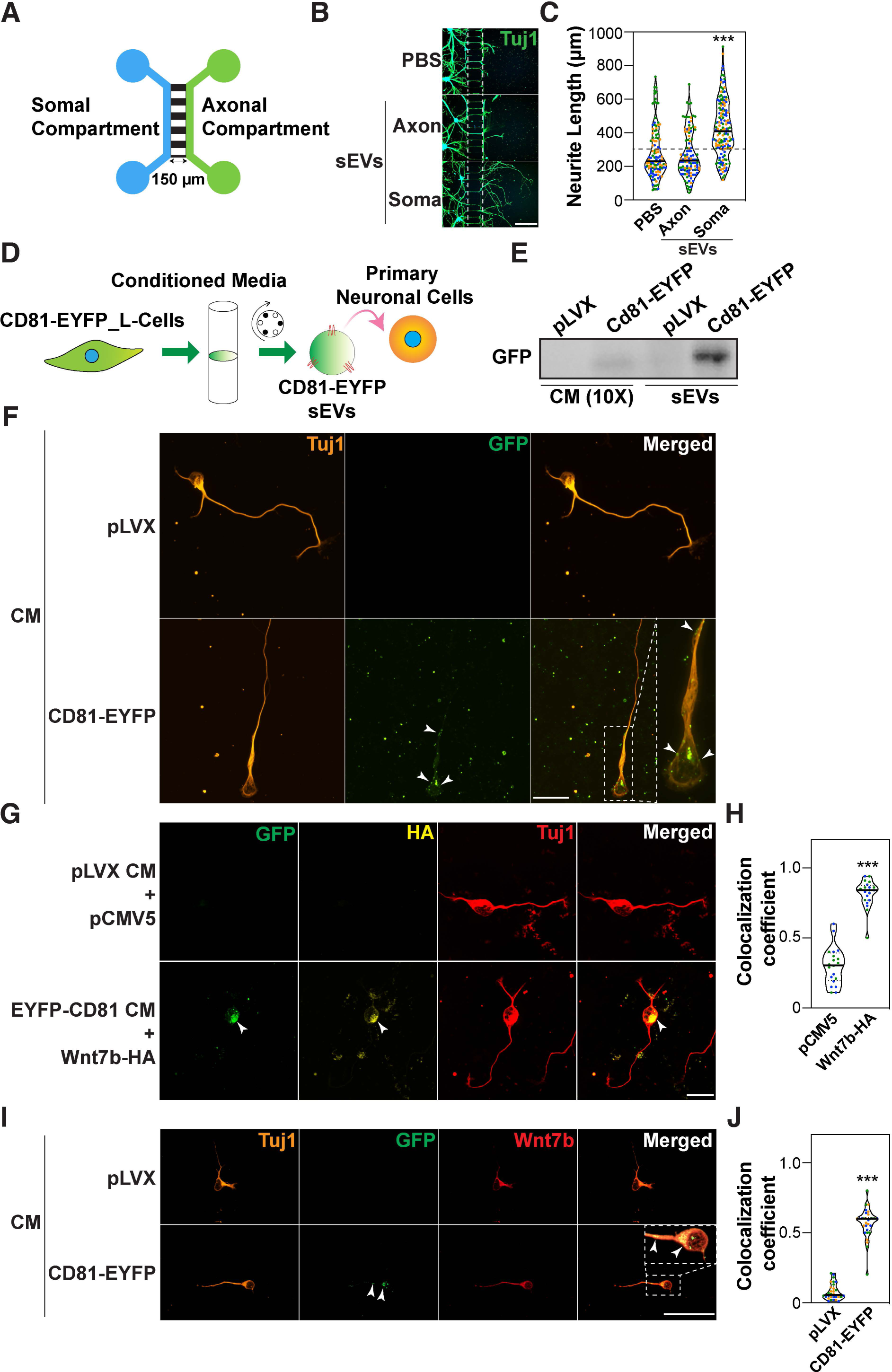
sEVs promote neurite elongation and co-localize with Wnt7b. **(A-C)** sEVs promote neurite elongation. **(A)** A schematic illustration of the two compartment Xona microfluidic device. The somal compartment is connected to axonal compartment through a 150 μm microgroove. **(B)** Cortical neurons (E15.5-16.5) were seeded in the somal compartment and cultured for 5 days prior to the addition of the L cell-derived sEVs in either the somal or axonal compartment. Neurons were fixed 24 h later and neuronal morphology was examined in Tuj1 stained neurons. Representative images are shown. Scale bar, 200 μm. **(C)** The length of the neurites growing in the microgroove and emerging in the axonal compartment was quantified for a minimum of 90 neurites. A dotted line marks both ends of the microgroove (150 μm). **(D-I)** sEVs can be internalized by neurons and associate with Wnts. Cortical neurons were treated with 10X concentrated conditioned media (CM) from L cells stably expressing CD81-EYFP, 4 h after plating for 29 h **(F)** or 24 h after plating for 30 min **(G, I)**. In panel **G**, cortical neurons were electroporated with 2 μg of either pCMV5 or C-terminal HA-tagged Wnt7b (Wnt7b-HA). Representative images of neurons immunostained with GFP and Tuj1 **(F, I)** or GFP, Tuj1 and HA **(G)** are shown. Scale bar, 20 μm **(F, G)** or 40 μm **(I)**. **(E)** Characterization of sEVs. The concentrated CM (10X) and sEV pellet from L cells were immunoblotted with anti-GFP antibody. **(H, J)** Pearson’s colocalization coefficient for **(G)** and **(I)**. Neurons were identified using Tuj1 as a reference channel and the colocalization coefficient was quantified using Nikon NIS-Elements software. Images **(F, I)** and the quantification **(J)** are representative of 30 neurons from 3 independent experiments or 20 neurons from 2 independent experiments **(G, H)**. In all the violin plots, values are distinctly colored for each experiment and the median marked by a black line. Statistical significance: ***p<0.001 using unpaired t-test **(H, J)** or one-way ANOVA with Dunnett’s post-test **(C)**.

To directly visualize sEV uptake, we used conditioned media (CM) from L cells stably expressing CD81-EYFP as a source of EYFP-labelled sEVs (Luga et al., 2012). Immunoblotting of both the conditioned media (CM) and the sEV pellet revealed the presence of EYFP using an anti-GFP antibody (Fig. 5D and 5E), indicating successful integration of the CD81-EYFP fusion protein in sEVs. Neurons were treated with the CM from either control or CD81-EYFP expressing L cells for 29 hours. Immunostaining followed by confocal microscopy revealed that the majority of GFP-positive neurons displayed abundant GFP puncta within the soma as well as occasional staining in the axon, suggesting that sEVs were internalized (Fig. 5F). Next, neurons were incubated with CM containing CD81-EYFP-labelled sEVs for 30 minutes, washed and subsequently chased in regular CM media for varying times prior to fixation. We found that sEVs were internalized by neurons within the 30 min exposure time, and that the internalized sEVs were found both in the soma and in axons throughout the time course (0 - 24 hours) (Fig. S8E). Thus, sEVs are rapidly (within 30 min) taken up by neurons and can be detected as long as 24 h later.

In cancer cells, internalized exosomes acquire endogenous Wnts to activate PCP signaling (Luga et al., 2012). Given that sEVs were internalized in neurons, we next investigated whether the internalized sEVs could associate with the Wnts. Cortical neurons were electroporated with HA-tagged Wnt7b (Wnt7b-HA) or empty vector control and 24 hours after plating were treated with EYFP-labelled sEVs for 30 min. Immunostaining with anti-HA and GFP antibody showed colocalization of Wnt7b-HA and sEVs in the soma (Fig. 5G and 5H). Next, we examined whether exosomes were colocalized with endogenous Wnt7b, the ligand most highly expressed in neurons. Neurons were treated with 10X concentrated CM obtained from either control or CD81-EYFP expressing L cells for 30 min and then were fixed and immunostained with anti-GFP and anti-Wnt7b antibodies, whose specificity was first confirmed by immunofluorescence microscopy (Fig. S8F). This analysis revealed colocalization of Wnt7b and sEVs in both the soma and axon (Fig. 5I and 5J). Given that internalized sEVs colocalize with Wnt7b, and that endogenous Wnts are required for sEV-induced outgrowth, our findings suggest that internalized sEVs may acquire neuronally expressed Wnt7b to promote axon outgrowth.

### sEVs from neurons and astrocytes do not promote neurite outgrowth

In the brain, exosomes mediate glia-neuron communication during developmental processes and pathogenesis (Chivet et al., 2013; Bahram Sangani et al., 2021; Gassama & Favereaux, 2021; Fan et al., 2022). Neurons and astrocytes are the two major cell types in the CNS, and are known to secrete exosomes (Gassama & Favereaux, 2021). Thus, we sought to determine whether exosomes released from neurons or astrocytes also modulate neurite outgrowth. To examine this, we first isolated and tested sEVs produced by primary embryonic cortical neurons (Fig. 6A). sEV pellets were positive for EV markers and displayed the typical 30-150 nm diameter size range (Fig. S9A and S9B). However, in contrast to fibroblast-derived sEVs, these sEVs did not promote neurite outgrowth when added to neurons at any dose tested, including at concentrations much higher (0.05-10 μg/mL) than those required for fibroblast-derived sEV effects (Fig. 6B and C). Thus, sEVs exogenously secreted from neurons do not promote neurite outgrowth.

**Figure 6.**
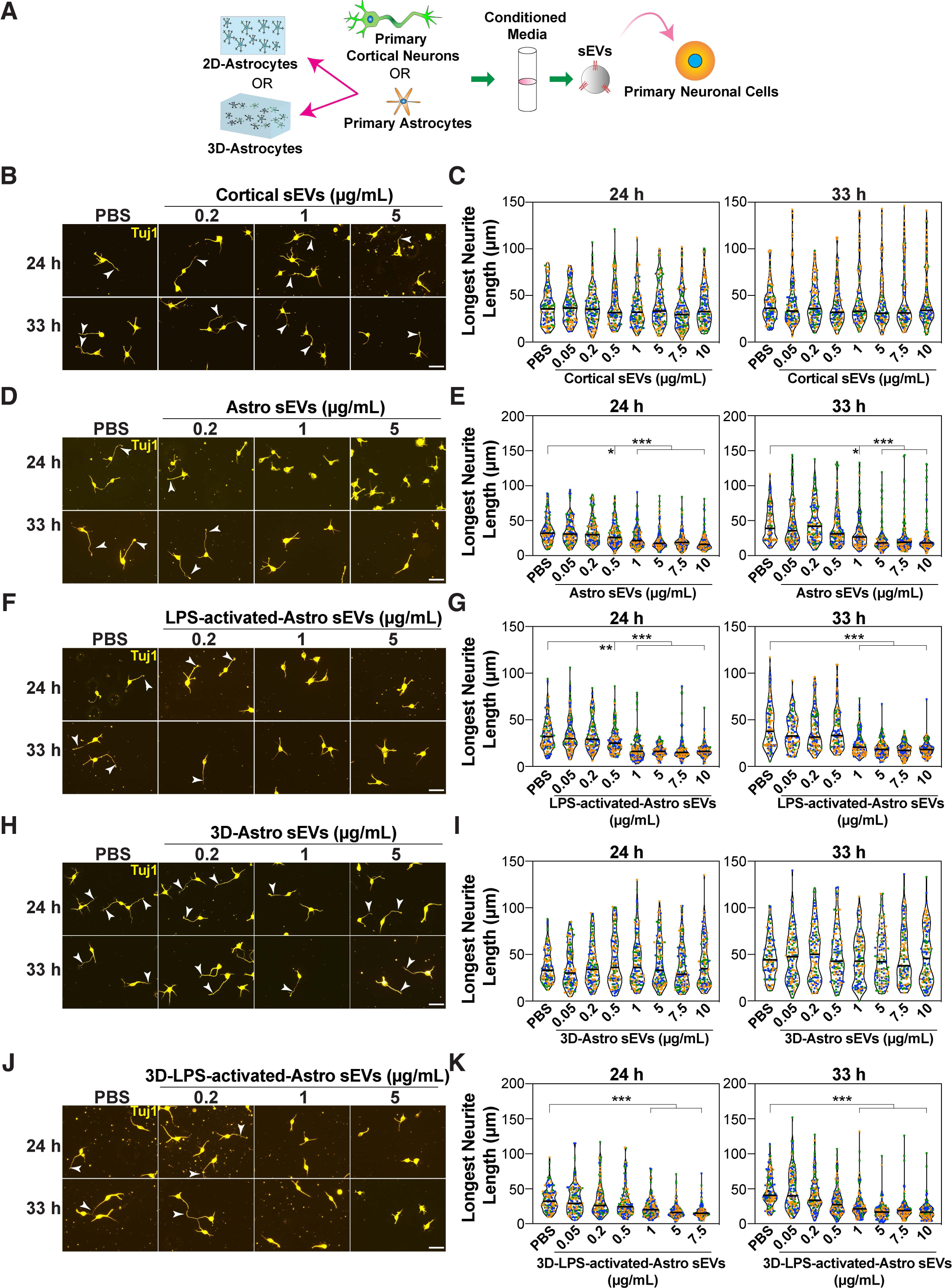
sEVs from neurons and astrocytes do not promote growth of the longest neurite. **(A)** A schematic illustration of the experimental set up. Dissociated E15.5-16.5 cortical neurons were treated with sEVs purified from primary cortical neurons or primary astrocytes. **(B-K)** Cortical neurons were treated with various concentrations (0.05-10 μg/mL) of sEVs purified from cortical neurons **(B, C)**, astrocytes **(D, E)**, LPS-activated astrocytes **(F, G)**, 3D-astrocytes grown in collagen gel **(H, I)** and LPS-activated 3D-astrocytes **(J, K)**, 4 h after plating. Neurons were fixed at 24 and 33 h, and neuronal morphology was examined in Tuj1 stained neurons. Representative images (**B, D, F, H, J**) and quantifications (**C, E, G, I, K**) are shown. Arrowheads mark the longest neurite. Scale bar, 40 μm. Neurite lengths are quantified from a minimum of 90 neurons per condition from 3 independent experiments and plotted as a violin plot with values from each experiment distinctly colored and the median marked by a black line. Statistical significance: *p<0.05, **p<0.01, ***p<0.001 using one-way ANOVA with Dunnett’s post-test.

Astrocytes support many physiological functions of neurons including synapse development and neuronal activity, whereas reactive astrocytes produced during injury or aging (Clarke et al., 2018) contribute to the production of the inhibitory environment that prevents axon regeneration (Clarke & Barres, 2013; Pekny & Pekna, 2014; Liddelow & Barres, 2017; Lee et al., 2022). Thus, we next investigated the role of astrocyte-derived exosomes on neurite outgrowth (Fig. 6A). For this, we isolated primary mouse astrocytes from newborn pups at postnatal day (P) 1-4 and confirmed their identity by positive staining for GFAP (Fig. S9C) and negative staining for the neuronal marker, Tuj1 (Fig. S9C). Reactive astrocytes were generated by treating isolated astrocytes with bacterial lipopolysaccharide (LPS), which induced upregulation of the expression of proinflammatory cytokines including TNF-α, iNOS-2, IL-1β and IL-6 as determined by qPCR (Fig. S9D), confirming astrocyte activation. sEVs were purified from control and LPS-activated astrocytes using differential centrifugation and the presence of sEVs in pellets confirmed by immunoblotting for CD81, TSG101 and Flotillin1 and by confirming particle diameters using NTA (Fig. S9E and S9F). Interestingly, treatment of neurons with varying doses of sEVs isolated from either control or LPS-activated astrocytes inhibited neurite outgrowth (Fig. 6D-G). As tissue culture plastic is much stiffer than the normal *in vivo* environment and can stimulate partial activation of astrocytes (East et al., 2009), we next cultured astrocytes in a soft 3D-collagen gel, hereafter referred as 3D-astrocytes. GFAP expression is higher in activated astrocytes (Escartin et al., 2021) and accordingly, astrocytes grown on a 2D-plastic surface showed much higher GFAP staining as compared to those cultured in the 3D-collagen gel (Fig. S9G). Moreover, treatment of 3D-astrocytes with LPS, enhanced GFAP expression (Fig. S9H). Next, sEVs were isolated from control and LPS-activated 3D-astrocytes and were confirmed to display typical exosome size using NTA (Fig. S9I-K). Remarkably, sEVs isolated from 3D-astrocytes had no effect on the neurite outgrowth (Fig. 6H and I), whereas sEVs from LPS-activated 3D-astrocytes inhibited the neurite outgrowth (Fig. 6J and K). Thus, our results indicate that while reactive astrocytes inhibit neurite outgrowth, control, unstimulated astrocytes grown in a soft environment do not alter neuronal morphology.

Taken together, these findings suggest that fibroblast-derived sEVs can promote neurite outgrowth, and this ability is not a general property of sEVs. Collectively, the data supports a model in which sEVs engage Wnt-PCP signaling in neurons to promote axon outgrowth and a polarized morphology. We suggest a novel mechanism whereby internalized sEVs which co-localize with Wnt ligands, such as Wnt7b, induce relocalization of the core PCP component, Vangl2 from the soma to distal part of the axon and thereby promote axon outgrowth (Fig. 7).

**Figure 7.**
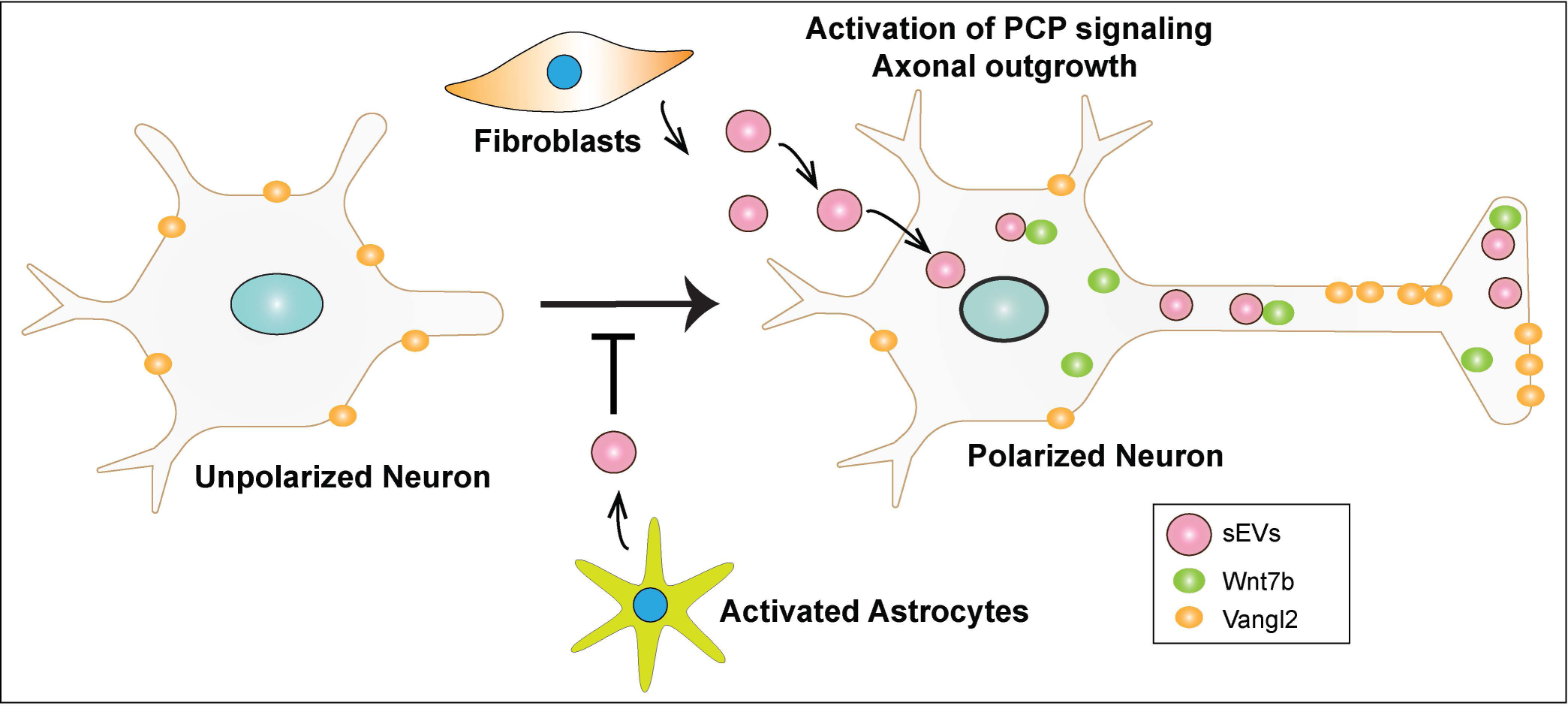
A model, depicting the effect of sEVs in promoting axon outgrowth and polarized neuronal morphology through Wnt-PCP signaling. sEVs secreted by L cells engage Wnt-PCP signaling in neurons to promote axon outgrowth that results in the acquisition of a polarized neuronal morphology. sEVs induce a shift in Vangl2 localization towards distal axon. sEVs can be internalized by neurons and can co-localize with Wnt7b to promote the growth of the prospective axon. In contrast to fibroblast derived sEVs, those isolated from activated astrocytes inhibit neurite outgrowth.

## Discussion

Exosomes are increasingly being studied for therapeutic applications (Dai et al., 2020; Kalluri & LeBleu, 2020). Exosomes from diverse sources are shown to modulate neurite outgrowth, but how fibroblast-derived exosomes act to specifically affect neuronal morphology remains unclear. Here, we show that fibroblast-derived sEVs can engage Wnt-PCP signaling in neurons and thereby promote axon outgrowth and acquisition of a polarized neuronal morphology. We demonstrate that sEVs are internalized via the soma in neurons and can colocalize with endogenous Wnt7b and that sEVs induce relocalization of the core PCP component, Vangl2 to the distal region of the axon. Thus, our work demonstrates that sEVs can engage Wnt-PCP signaling in neurons to promote axon outgrowth.

Exosomes derived from diverse sources harbour the ability to promote axon or dendrite elongation. For instance, exosomes from Schwann cells of the PNS have been shown to promote neurite outgrowth in dorsal root ganglia (DRG) neurons, and exosomes from mesenchymal cells enhance neurite outgrowth in DRG and CNS-derived cortical neurons (Xin et al., 2012; Lopez-Verrilli et al., 2013; Lopez-Verrilli et al., 2016; Zhang et al., 2017). In the case of fibroblasts, exosomes from these cells can promote neurite outgrowth in DRG neurons in normal, uninjured contexts, and can enhance axon regeneration in injured sciatic nerves (Zhao et al., 2023). In addition, fibroblasts can promote Schwann cell migration and sorting, both of which are associated with axonal regrowth (Dreesmann et al., 2009; Parrinello et al., 2010; van Neerven et al., 2013). In our study, we found that fibroblast-derived sEVs can promote axon outgrowth in cortical neurons and that they act through the Wnt-PCP pathway. In contrast, in a prior study, fibroblast-derived exosomes were shown to promote axon regeneration in a Wnt-dependent manner but did so via the mTOR pathway and only in neurons grown in an inhibitory environment (Tassew et al., 2017). Thus, exosomes appear to be capable of activating distinct neuronal signaling pathways in growth permissive versus inhibitory contexts. In contrast to fibroblasts, we observed that sEVs derived from primary neurons or unactivated astrocytes had no effect on neurite outgrowth even when used at higher concentrations. Upon damage, activated astrocytes contribute to the inhibitory microenvironment by forming a glial scar at the injury site (Pekny & Pekna, 2014; Liddelow & Barres, 2017; Lee et al., 2022). In line with this, we observed that sEVs derived from LPS-activated astrocytes inhibit neurite outgrowth, which is consistent with a previous report showing that exosomes from Interleukin-1β-activated human primary astrocytes similarly inhibited neurite outgrowth in mouse cortical neurons (You et al., 2020). This suggests that the cell type and state of exosome secreting cells determines the axon growth promoting properties of exosomes. Understanding the mechanisms that alter exosome cargos to thereby modulate the effects on neurons is an important area for future investigation.

Core PCP components modulate many aspects of neuronal morphology including neuronal polarization, axon outgrowth, axon guidance and synapse formation (Goodrich, 2008; Davey & Moens, 2017; Hakanen et al., 2019; Zou, 2020). Our data shows that both of the related PCP components, Pk1 and Pk2 are required for growth of the prospective axon, which is consistent with a recent study showing that Pk2 is necessary for proper neuronal polarity in rat hippocampal neurons (Dorrego-Rivas et al., 2022). We also observed that the absence of Pk1 and Pk2 attenuated sEV-induced growth of the longest neurite. Another core PCP component, Vangl2 has been reported to exhibit differential effects in distinct neurons. For instance, downregulation of Vangl2 in commissural neurons using shRNAs inhibited Wnt5a-induced axonal outgrowth (Shafer et al., 2011), whereas hippocampal neurons from Vangl2 conditional-knockout mice exhibited increased basal and Wnt5a-induced axon outgrowth (Dos-Santos Carvalho et al., 2020). In our study, we observed that both Vangl1 and Vangl2 are required for basal neurite outgrowth and that only Vangl2 is required for sEV-induced neurite outgrowth. Importantly, sEVs induced relocalization of Vangl2 to the distal part of the axon, further supporting a role for Vangl2 in sEV-mediated axon outgrowth. Although not designated as core PCP components, members of the Smurf E3 ubiquitin ligase family are important regulators of PCP, and can modulate neuronal morphology (Vohra et al., 2007; Narimatsu et al., 2009; Cheng et al., 2011; Kannan et al., 2012). Consistently, we found that both Smurf1 and Smurf2 are required for neurite outgrowth, and that sEV-induced neurite outgrowth is attenuated in Smurf1 and Smurf2 knockout neurons. Taken together, these results demonstrate that sEVs engage key components of the Wnt/PCP pathway to modulate neuritogenesis.

Fzd receptors and Dvls are essential for PCP signaling, but also mediate canonical Wnt/μ-catenin signals (van Amerongen et al., 2008). Diverse Fzd receptors have been shown to modulate events that establish neuronal morphology including axonal polarity and growth, axon guidance, dendrite morphogenesis and synaptogenesis (Pascual-Vargas & Salinas, 2021). In particular, Fzd3, 5 and 9 have been reported to be essential for axon outgrowth and guidance (Pascual-Vargas & Salinas, 2021). In embryonic cortical neurons (E15.5-16.5), we observed that Fzd1, 2, 3, 7, 8 and 9 are expressed at varying levels. Of these Fzds, individual knockdown of Fzd1, 3 or 9 attenuated both basal and sEV-induced neurite outgrowth, whereas Fzd7 was only required for neurite outgrowth mediated by sEVs. This indicates that Fzd1, 3 and 9 act in a non-redundant manner and that Fzd7 is specifically required for sEV-mediated effects. Consistently, a non-redundant role for Fzd3 in mediating axon outgrowth and guidance in knockout mouse models has been described (Hua et al., 2013; Hua et al., 2014; Qu et al., 2014). The cell surface and intracellular distribution of Fzds can be modulated by post-translational modifications such as phosphorylation (Pascual-Vargas & Salinas, 2021). Moreover, in neurons, a distinctive spatiotemporal localization of Fzds in the soma, axon and growth cone during different developmental stages has been observed (Sahores et al., 2010; Varela-Nallar et al., 2012; Slater et al., 2013). This differential localization of Fzds in space and time, due at least in part to changes in post-translational modifications, could be critical for axon formation and outgrowth, and thus may explain the non-redundant role of Fzds during this process. Dvls are shared by PCP and canonical Wnt pathways and have been reported to modulate axon outgrowth and guidance (Gao & Chen, 2010; Sharma et al., 2018). For instance, Dvl2 has been reported to specifically function in PCP signaling (Onishi et al., 2013). We observed that cortical neurons express all three Dvls (Dvl1, 2 and 3), but only Dvl2 was required for sEV-induced neurite outgrowth, a process that we have shown requires PCP signaling. Altogether our data indicates a key role for the PCP pathway in regulating neuronal polarization and axon outgrowth that is enhanced by fibroblast-derived sEVs.

In vertebrates, Wnts are required for the activation of PCP signaling and can modulate neuronal morphology (Butler & Wallingford, 2017; Teo & Salinas, 2021; Koca et al., 2022). Our data shows that neuronal Wnt secretion is required for axon formation as chemical inhibition of porcupine or siRNA-mediated knockdown of Wls, both of which block Wnt secretion, prevented axonogenesis. This observation is consistent with a previous report showing that treatment with the porcupine inhibitor, IWP2, inhibits neuronal polarization in hippocampal neurons (Stanganello et al., 2019). Moreover, we observed that sEV-induced neurite outgrowth was inhibited upon blocking neuronal Wnt secretion. We showed that cortical neurons express Wnt5a, 5b, 7a and 7b, three of which (Wnt5b, 7a and 7b) are required for the growth of the prospective axon and for enhanced axon elongation induced by sEVs. Intracellular association of Wnts with exosomes is critical for Wnt secretion on exosomes (Routledge & Scholpp, 2019; Gross, 2021; Mittermeier & Virshup, 2022). Consistent with this, we showed that neuronal Wnt7b, the most highly expressed Wnt in E15.5 cortical neurons, can colocalize with the exogenously applied CD81-positive sEVs within the neurons. This suggests the possibility that sEVs internalized by neurons can engage endogenous Wnts in the context of neurite outgrowth. In a prior study in cancer cells, we showed that exosomes from fibroblasts acquire Wnt11 when internalized by breast cancer cells to activate autocrine Wnt-PCP signaling and promote cell motility and formation of protrusions (Luga et al., 2012). Thus, we speculate that sEVs might similarly activate autocrine Wnt-PCP signaling in neurons to promote axon outgrowth, though precisely how, remains to be explored.

Although the fibroblasts used in our study were derived from skin, lung and mouse embryos, and thus do not reside in CNS, the current study identifies a potential of fibroblast-derived sEVs, irrespective of the tissue origin, in activating axon outgrowth systems, and thus provides evidence of the therapeutic potential of fibroblast sEVs in treating neuronal injury. At the molecular level, we have shown that sEVs induce axon elongation and a polarized neuronal morphology by engaging Wnt-PCP signaling components. As context-dependent effects of exosomes are not well understood, it will be critical to explore the molecular determinants in exosomes that confer context specificity in varying biological functions.

## Acknowledgements

We thank Lisa Qiu, Victoria Leon Guerrero, Bruno Sangiorgi and Nadia Zafar for contributions during the early stages of the project. We thank Dr. Sachdev Sidhu for kindly providing Fzd blocking antibody. This work was funded by grants to L.A. from the CIHR (FDN148455) and to L.A. and J.W. from the Krembil Foundation.

## Materials and Methods

### Mice

Mouse studies were conducted in accordance with the guidelines of the Canadian Council on Animal Care (CCAC) and approved by the University Animal Care Committee, University of Toronto or The Centre for Phenogenomics (TCP), Toronto, Ontario, Canada. Mice were housed in individually ventilated cages with free access to standard animal food and water in a climate-controlled room with a 12 h light/dark cycle. CD1 pregnant females E15-16 were purchased from Charles River.

Prickle1 (Pk1) conditional knockout mice were generated by introducing Pk1 targeting vector (EUCOMM, HTGR03009_Z_6_E08) into G4 embryonic stem (ES) cells by electroporation. ES cell colonies were selected using Neomycin (Invitrogen, 10131027) and genomic DNA was analyzed by southern blotting to identify homologous recombinants. Chimeric embryos were produced, at the Transgenic Core of the TCP facility, by diploid aggregation of in house generated Pk1 or commercially acquired targeted Pk2 (HEPD0638_2_E01 clone C3) ES cells and CD1 embryos. Chimeric mice were crossed with ICRs mice, obtained from TCP, to obtain the F1 heterozygotes, that were then crossed with Flp recombinase mice to remove the neomycin selection cassette. For the Pk1 allele, genomic DNA was digested with NdeI and probed with a Pk1 3′ probe, resulting in a 15.7 kb (wild-type allele) and a 10.4 kb (mutant allele) fragment. For the Pk2 allele, genomic DNA was digested with BstEII, ScaI, NsiI, AvrII, or Bsu36 and probed with a Neo probe to produce targeted bands of 15Kb, 18.5Kb, 17.6 Kb, 9.5Kb or 12.7Kb, respectively. Single floxed mice (Pk1^fl/fl^; Pk2^+/+^ and Pk1^+/+^; Pk2^fl/fl^) were crossed with each other to obtain Pk1/2 double floxed mice (Pk1^fl/fl^; Pk2^fl/fl^). To obtain the brain specific conditional knockout, single floxed mice and double floxed mice were crossed with a Nestin-Cre line, acquired from TCP, to obtain Cre+ (Pk1^fl/fl^;Pk2^+/+^;Cre+, Pk1^+/+^;Pk2^fl/fl^;Cre+, Pk1^fl/fl^;Pk2^fl/fl^;Cre+) and Cre-(Pk1^fl/fl^; Pk2^+/+^;Cre-, Pk1^+/+^;Pk2^fl/fl^;Cre-, Pk1^fl/fl^; Pk2^fl/fl^;Cre-) floxed mice that were subsequently crossed with each other to obtain the various genotype combinations. For genotyping, genomic DNA from 2 mm E16.5 tail piece was isolated and then subjected to PCR using primers listed in Table S1. Loss of Pk1/2 expression was confirmed in brain sections from Pk1^fl/fl^; Pk2^fl/fl^;Nestin-Cre+ mice by immunostaining with Pk1/2 antibodies (Table S3), followed by confocal microscopy.

Heterozygous Vangl2^Lp/+^ (Vangl2^S464N^) mice (Strong & Hollander, 1949), obtained from Jackson Labs (#000220), were kindly provided by Dr. Helen McNeil (Lunenfeld-Tanenbaum Research Institute, Mount Sinai Hospital, Toronto, Ontario, Canada) and crossed with each other to obtain homozygous mutants (Vangl2^Lp/Lp^). Smurf1/2 heterozygous mice (Smurf1^+/−^; Smurf2^+/−^) were generated previously (Narimatsu et al., 2009), and crossed with each other to generate single knockout (Smurf1^−/−^ or Smurf2^−/−^) embryos.

### Isolation, culturing and transfection of mouse cortical neurons

Primary cortical neurons were isolated from mouse embryos E15.5-16.5 as previously described (Christova et al., 2023). Briefly, brains were harvested and placed in ice-cold HBSS (Gibco) and cerebral cortices were dissected. Cortices were digested in 0.25 % Trypsin-EDTA (Gibco, 25200056) at 37°C followed by trypsin inactivation by adding Dulbecco’s Modified Eagle Medium (DMEM) (Gibco, 11995-065) containing 10% FBS (Gibco, 12483-020). After further mechanical dissociation, neurons were seeded at a density of 30000 cells/well for regular experiments or 125,000 cells/well for transfection experiments in 4-well chamber slides (Lab-Tek II, 155382) coated with poly-L-lysine (20 µg/mL, Sigma, P1399) and laminin (2 µg/mL, Corning, 354232). Neurons were cultured at 37°C and 5% CO_2_ in Neurobasal medium (Gibco, 21103-049) supplemented with 2% B-27 (Gibco, 17504-044), 0.5% N-2 (Gibco, 17502-048), 2 mM GlutaMAX^TM^ (Gibco, 35050-061) and 0.5% penicillin/streptomycin (Gibco, 15140-122).

For transfections, dissociated neurons were electroporated with cDNA (2 µg) or siRNA (Table S2) using an Amaxa nucleofector (program-005, Lonza) and mouse neuron nucleofector^TM^ kit (VPG-1001, Lonza) according to the manufacturer’s instructions. Briefly, neurons (4-5 × 10^6^) were electroporated with a mix of 2 µM siGENOME SMARTpool siRNAs (Dharmacon), comprised of a pool of four individual siRNAs except for siWnt7a pool which contained two individual siRNAs (Table S2), and 2 µg of enhanced green fluorescent protein (eGFP) as a transfection reporter. Media was replaced 4 h after seeding. All DNAs were prepared using Endotoxin-Free Maxi-prep kit (Qiagen).

### Mammalian cell culture and transfection

L cells (CRL-2648), Human Dermal fibroblasts (HDFn) (PCS-201-010™), BJ cells (CRL-2522^TM^) and MDA-MB-231 cells were purchased from ATCC, normal Human Lung Fibroblasts (NHLF) were purchased from Lonza (CC-2512) and mouse embryonic fibroblasts (MEFs) were acquired from the TCP facility. MDA-MB-231 cells were cultured in Roswell Park Memorial Institute (RPMI) 1640 medium containing 5% FBS while all other lines were cultured in DMEM containing 10% FBS. Cell lines were routinely tested for mycoplasma contamination using MycoAlert^TM^ Plus reagent (Lonza, LT07-703).

For siRNA transfections, L cells were (1 × 10^6^) were transfected with 40 nM siGENOME SMARTpool siRNAs (Dharmacon) using Lipofectamine^TM^ RNAiMAX reagent (Invitrogen, 13778075) according to manufacturer’s protocols. For cDNA transfection of MDA-MB-231 cells, plasmids (2 µg) were delivered using Lipofectamine^®^ 3000 (Invitrogen, L3000-001) according to manufacturer’s instructions. The Wnt7b construct was generated using Gateway system (Life Technologies) by transfer from an entry clone into a C-terminal HA-tagged pCMV5-based destination vector.

### Primary Astrocyte Cultures

Dissociated primary astrocytes were isolated from the cortices of CD1 mouse post-natal (P1-P4) pups as previously published (Schildge et al., 2013). Dissociated cells were cultured in DMEM supplemented with heat inactivated 10% FBS and 1% penicillin/streptomycin in 75 cm^2^ flasks (Corning, 430720U) coated with poly-D-lysine (50 µg/mL, Sigma, P7280). Media was changed after 24 h of initial seeding, and every 2 days thereafter. Cultures were grown for 8 days. and then shaken at 120 rpm overnight to remove microglia and less adherent cells. Cells were then washed, trypsinized and these astrocyte enriched cultures were used for subsequent experiments.

For 3D cultures, astrocytes were seeded in a gel containing type-I rat tail collagen (Ibidi, 50201), prepared according to the manufacturer’s protocol. Briefly, 10% minimum essential media (MEM, Gibco, 11430-030), 1% GlutaMAX^TM^ and collagen (2 mg/mL) were mixed on ice and neutralized using 1M NaOH. Subsequently, DMEM containing cells was gently added to the gel mixture resulting in a cell density of 2 × 10^6^ cells/mL of gel mixture. Gel mixture was transferred to either 6-well plates (2.5 mL/well) for sEV isolation or in chamber slides (700 μL/well) for immunofluorescence microscopy. The liquid gel mixture was placed at 37°C and 5% CO_2_ for 30 minutes to solidify and then fresh DMEM containing heat inactivated 10% FBS and 1% penicillin/streptomycin was added.

### Preparation of Conditioned Media

Fibroblasts (0.5 × 10^6^) were seeded in a 100 mm dish and grown to confluency. Cells were washed with DMEM and incubated with DMEM either without FBS (L cells) or exosome free 10% FBS (for NHLF, HDFn, BJ, MEFs) (Gibco). After 3-4 days, conditioned media (CM) was collected, centrifuged at 300 x g to remove cells and stored at 4 °C. CM from neurons was prepared from primary cortical neurons (4 × 10^6^ cells) seeded in a 100 mm dish and cultured in a serum free complete medium. On day 4, 25% of freshly made complete medium supplemented with 5 µM cytosine β-d-arabinofuranoside (Sigma, C1768) was added and CM was collected on day 8. For 2D astrocytes, enriched primary astrocytes cells were seeded in p75 flasks and cultured in DMEM supplemented with heat inactivated exosome free 10% FBS. Cultures were grown to confluency, and the CM was collected after 6 days. For 3D astrocytes cultures in collagen gel, enriched primary astrocytes were seeded in 6 well plate and CM was collected after 6 days. Small EVs from fibroblast-CM were purified within 1 week, whereas sEVs from neurons and astrocytes were isolated on the same day.

### Isolation of small extracellular vesicles (sEVs)

Conditioned media (CM) (50 mL) was collected from fibroblasts, neurons or astrocytes and centrifuged at 300 x g for 5 min and at 2000 x g for 20 min to remove cells and large debris, respectively. Supernatants were then centrifuged at 10,000 x g for 30 min to remove apoptotic bodies and large vesicles. The resulting supernatants were filtered through a 0.2 μm filter (Thermo Fisher Scientific, 568-0020) and ultracentrifuged at 100,000 x g (Type 70 Ti rotor, Beckman Coulter) for 2 h using 25 ξ 89 mm polycarbonate tubes (Beckman Coulter, 355618). Pellets were resuspended in 500 μL of sterile phosphate-buffered saline (PBS) and diluted with 7.5 mL of PBS. Diluted pellets were ultracentrifuged at 100,000 x g (Type 70.1 Ti rotor, Beckman Coulter) for 2 h using 16 ξ 76 mm polycarbonate tubes (Beckman Coulter, 355603). The resulting pellets were resuspended in 250 μL of sterile PBS, aliquoted and stored at −80°C.

### Purification of sEVs using iodixanol density gradient

A discontinuous iodixanol gradient, prepared by diluting 60% iodixanol (Optiprep^TM^, Sigma, D1556) with 0.25 M sucrose / 10 mM Tris, pH 7.4, was used to further purify the sEV pellets (100,000 x g). The sEV pellet, resuspended in 500 μL of PBS, was overlaid on top of the discontinuous gradient (40%, 20%, 10% and 5%; 3 mL each, but 2.5 mL for 5%) and ultracentrifuged at 100,000 x g (SW 40 Ti or SW41 Ti rotor, Beckman Coulter) for 16 h using ultra-clear tubes (Beckman Coulter, 344060 or 344059). Twelve fractions of 1 mL each were collected manually from the top. Fractions were washed by diluting with 7 mL of PBS and ultracentrifuged at 100,000 x g (Type 70.1 Ti rotor, Beckman Coulter) for 2 h. The resulting pellets were resuspended in 150 μL of sterile PBS and stored at −80°C.

### Purification of sEVs by size exclusion chromatography

The sEV pellets (100,000 x g) were purified using qEV original size exclusion column with a pore size of 70 nm (Izon science, 1001683) according to manufacturer’s protocol with minor modifications. Briefly, the column was equilibrated with 30 mL of PBS and exosome pellet, resuspended in 500 μL of PBS, was overlaid on qEV column followed by elution with PBS. Fractions of 500 μL were collected and analyzed for particle and protein concentration using NTA and Bradford assay, respectively.

### Purification of sEVs by Exo-spin^TM^ precipitation

The sEVs from 3D-astrocytes were purified using Exo-spin^TM^ purification kit (Cell Guidance Systems) according to manufacturer’s instructions. Briefly, CM was precleared by sequential centrifugations at 300 x g, 2000 x g and 10,000 x g. Supernatants were mixed with the precipitation buffer and incubated overnight at 4°C. Samples were centrifuged at 10,000 x g for 30 min and pellets were re-suspended in 200 μL of PBS. Samples were further purified using provided size exclusion columns and exosomes were eluted in 200 μL of PBS.

### Proteinase K and detergent treatment of sEVs

sEVs were incubated with a final concentration of 100 μg/mL Proteinase K or 1% Triton X-100 at 37°C for 1 hour with gentle vortexing every 15 min. After treatment, sEVs were washed in 7 mL of PBS, ultracentrifuged for 2 h and the resulting pellet was resuspended in 120 μL of PBS.

### Nanoparticle Tracking Analysis

The number of particles were measured using ZetaView^®^ PMX 110 device (Particle Metrix GmbH, Germany). The measurements were taken at all 11 different positions with camera sensitivity set to 90 and shutter speed at 45 using the ZetaView software (version 8.03.08). The video quality was set to medium, and the data was analyzed using the ZetaView analysis software (version 8.02.31) with the following post-acquisition parameters: minimum size 10, maximum size 1000, and minimum brightness 20.

### Small EV internalization assay

For sEV internalization assays, 50 mL of media was collected from L cells stably transfected with either CD81-EYFP or CD81-pLVX as a control (Luga et al., 2012). CM was centrifuged at 300 x g for 5 min and at 2000 x g for 20 min to remove cells and large debris, respectively. Supernatants were concentrated 10-fold using a 10 kDa MWCO filter (Amicon® Ultra-15, UFC901008). Neurons were seeded in 8 well Ibidi chambers and treated with 10X CM for 29 h or 30 min, prior to fixation, immunostaining and imaging using confocal microscopy.

### Neurite outgrowth assay in a microfluidic chamber

Two compartment microfluidic chambers containing a 150 μm microgroove were purchased from Xona Microfluidics (XC150). Chambers were prepared for seeding neurons according to manufacturer’s instructions with a few modifications. Briefly, chambers were treated with a pre-coat solution and subsequently coated with poly-L-lysine (20 µg/mL) and laminin (2 µg/mL). Cortical neurons (50,000 cells) were seeded in one well in an initial volume of 20 μL and allowed to adhere for 5 min at 37°C and 5% CO_2_. Complete Neurobasal medium was added to all the wells and neurons cultured for 5 days until axons crossed the microgroove and emerged in the distal chamber. The sEVs (5 μg/mL) were added either to the somal side or axonal sides on day 5 for 24 h prior to fixation and immunostaining.

### RT and qPCR

For neurons and astrocytes, total RNA was extracted using the Norgen Single-Cell RNA Purification Kit (Norgen, 51800). For other cells, total RNA was extracted using PureLink RNA Mini Kit (Life Technologies, 12183025). Complementary DNA was synthesized using 35 to 200 ng of purified RNA using oligo-dT primers and M-MLV Reverse Transcriptase (Invitrogen, 28025-013). Real-time qPCR was performed using SYBR^TM^ Green PCR Master Mix (Applied Biosciences, 4309155) on the QuantStudio 6 Flex System (Applied Biosciences). Relative gene expression was quantified using the ΔΔCt method and normalized to glyceraldehyde-3-phosphate dehydrogenase (Gapdh). Primers used are listed in the Supplementary Table S4.

### Immunofluorescence microscopy and immunoblotting

For immunocytochemistry, neurons and astrocytes were fixed in 4% paraformaldehyde as previously described (Christova et al., 2023). Briefly, cells were permeabilized with 0.2% Triton/PBS, washed with PBS, and blocked at room temperature for 1 h with 3% Bovine Serum Albumin (BSA)/PBS. Cells were incubated with primary antibodies for 2 h at room temperature or overnight at 4°C, washed 3 times with PBS, and incubated with secondary antibodies for 2 h at room temperature. Cells were washed with PBS and stored in PBS or in Ibidi mounting medium prior to imaging. For immunohistochemistry of mouse embryos at E15.5, embryos were fixed in 4% paraformaldehyde in PBS at 4°C overnight. After washing with 0.1% Tween 20 in PBS (PBST) 3 times for 5 min, the samples were cryoprotected by immersing in 15% sucrose in PBS at 4°C overnight, then 30% sucrose in PBS at 4°C overnight. Samples were embedded in optimum cutting temperature (OCT) compound and cyrosections of 16 μm were prepared with a cryostat (Leica Biosytems, CM3050 S), which were then stained with antibodies listed in Table S3. For epifluorescence microscopy, images were acquired using the 40X Plan-NEOFLUAR objective of Zeiss Axiovert 200 M epifluorescence microscope. For confocal microscopy, images were acquired using Zeiss AXIO Observer Z1 microscope equipped with Zen Blue software, a spinning disc confocal scanner (CSU-X1, Yokogawa) and a CCD camera (Axiocam 506 mono) with following objectives: Plan-Apochromat 20x/0.8 M27 air, Plan-Apochromat 40x/1.4 Oil DIC and Plan-Apochromat 63x/1.4 Oil DIC M27. Some confocal images were acquired using Nikon ECLIPSE Ti2 inverted microscope equipped with LUA-S laser unit and NIS-Elements software (from Nikon) with following objectives: Nikon-N Apo-LWD 40X/1.15 WI and Nikon-N Apo-LWD 20X/0.95 WI. All images were acquired in a random fashion.

For immunoblotting, cells were lysed in lysis buffer (50 mM Tris-HCl, 150 mM NaCl, 1 mM EDTA, 0.5% Triton X-100, 1 mM DTT containing protease and phosphatase inhibitors) (Bao et al., 2012). Protein concentration was determined using Bradford reagent (BIORAD). Proteins from cell lysates (1-20 μg) and exosomes (1-10 μg) were separated on an SDS-PAGE gel and analyzed by immunoblotting as described previously (Heidary Arash et al., 2014).

### Electron microscopy

Exosomes were visualized using Talos L120C transmission electron microscope (TEM) (Thermo Scientific) in the Microscopy Imaging Laboratory, University of Toronto, Canada. The sEV samples were frozen on TEM grids (Quantifoil R2,2 300 mesh, EMS) that were charged for 30 seconds with a Pelco EasiGlow glow discharge cleaning system (Ted Pella). Samples (4 μL) were deposited onto the grid and the grid was blotted and plunge frozen using a Vitrobot IV (FEI). Images were taken using 120kV at magnifications 28000x and 57000x, corresponding to a pixel size of 510 and 249 pm respectively. Images were collected at a defocus of −4 to −1.5 microns using the TEM Image and Analysis (TIA) software.

### Quantification

For Tuj1 stained neurons, the length of all neurites (including the longest neurite, a prospective axon, and dendrites, all other neurites excluding the longest neurite) were quantified using Volocity software (Quorum Technologies Inc, Canada). In MAP2 and Tau-1 experiments, neurons were classified into two groups: only MAP2-positive minor neurites or minor neurites and a single Tau-1-positive axon displaying medial-to-distal localization of the Tau-1.

For Vangl2 localization, the intensity of Vangl2 in soma, proximal axon and distal axon was quantified using Nikon NIS-Elements software (Version 5.41.00). For sEVs and Wnt7b co-localization, neurons were identified using Tuj1 as a reference channel and the Pearson’s colocalization coefficient for GFP (sEVs) and Wnt7b was determined using Nikon NIS-Elements software (Version 5.41.00).

### Statistical Analyses

The data for the axon length and longest neurite length is plotted as a violin plot from 90-120 neurons per condition in 3 independent experiments. The data was analyzed statistically using an unpaired t-test for comparing two groups or one-way ANOVA with Dunnetts post-test for multiple comparisons to a single control group, or two-way ANOVA with Tukey’s post-test for multiple comparisons to control and between samples, as indicated in the figure legends. All statistics were done in GraphPad PRISM 9 (GraphPad software Inc. La Jolla, CA). Statistical significance: *p<0.05, **p<0.01, ***p<0.001.

---

**Figure S1.**
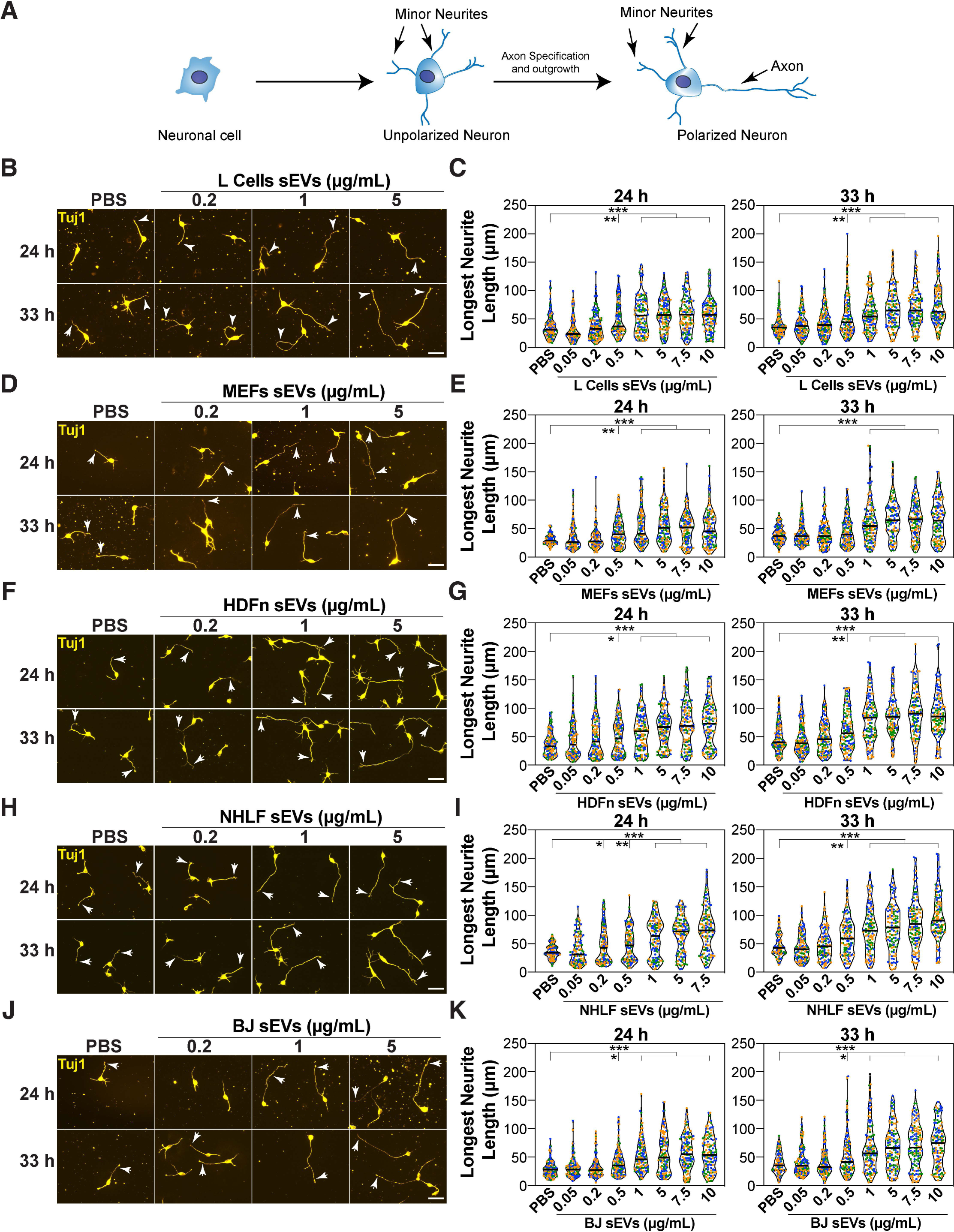
sEVs promote the growth of the longest neurite in a concentration-dependent manner. Related to Fig. 1. **(A)** A schematic of the morphological changes in neurons cultured *in vitro*. **(B-K)** Dissociated E15.5-16.5 mouse cortical neurons were treated with various concentrations (0.05-10 μg/mL) of sEVs purified from the indicated fibroblast cell lines including L cells **(B, C)**, MEFs **(D, E)**, HDFn **(F, G)**, NHLF **(H, I)** and BJ **(J, K),** 4 h after plating. Neurons were fixed at 24 and 33 h, and neuronal morphology was examined in Tuj1 stained neurons. Representative images **(B, D, F, H, J)** and quantifications **(C, E, G, I, K)** are shown. Arrowheads mark the longest neurite. Scale bar, 40 μm. Neurite lengths are quantified from a minimum of 90 neurons per condition from 3 independent experiments and plotted as a violin plot with values from each experiment distinctly colored and the median marked by a black line. Statistical significance: *p<0.05, **p<0.01, ***p<0.001 using one-way ANOVA with Dunnett’s post-test.

**Figure S2.**
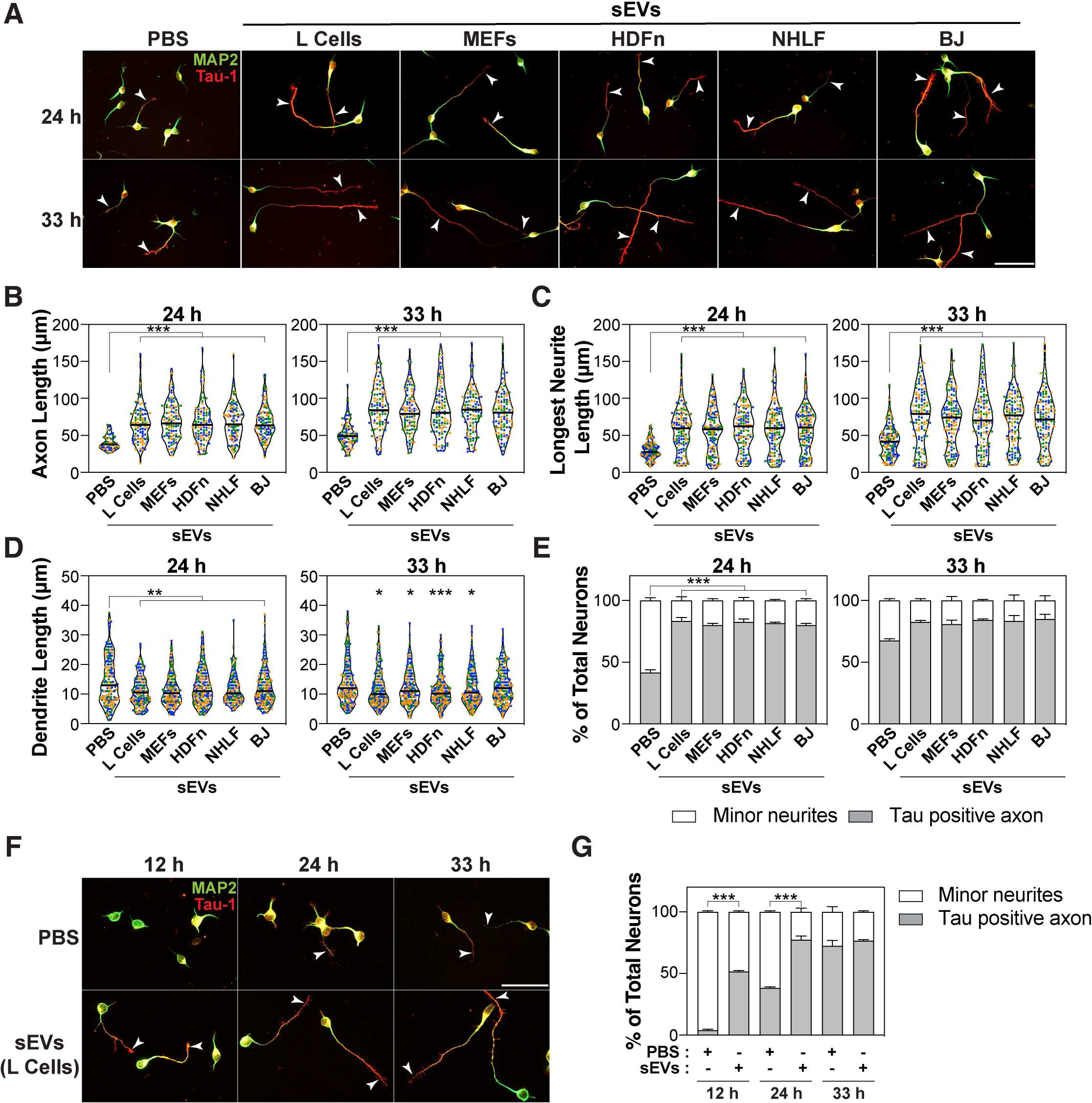
sEVs promote axon outgrowth and polarized neuronal morphology. Related to Fig. 1. (**A-G)** Dissociated E15.5-16.5 mouse cortical neurons were treated with sEVs purified from the indicated fibroblast cell lines **(A-E)** or only L cells **(F-G)**, 4 h after plating. Neurons were fixed at 24 and 33 h **(A-E)** or 12, 24 and 33 h **(F-G)**, and neuronal morphology was examined in neurons immunostained for MAP2 (dendrites, green) and Tau-1 (axons, red). **(A, F)** Representative images are shown. Arrowheads mark Tau-1 positive axons. Scale bar, 50 μm. (**B-E**) The length of the Tau-1 positive axons **(B)**, longest neurite (independent of Tau-1+ staining, **C**) and individual dendrite length (MAP2 positive, **D**) is quantified. (**E, G)** Percent of neurons containing only minor neurites (MAP2 positive) or a single Tau-1-positive axon, is plotted. Neurite lengths are quantified from a minimum of 45 (for 24 h in panel **B**) or 90 (for all others) neurons per condition from 3 independent experiments and plotted as a violin plot with values from each experiment distinctly colored and the median marked by a black line. In panel **B**, for the 24 h control group (PBS treated), less data points were available due to greater percentage of neurons lacking a Tau-1 positive axon. Statistical significance: *p<0.05, **p<0.01, ***p<0.001 using one-way ANOVA with Dunnett’s post-test **(B, C, D)**, or two-way ANOVA with Tukey’s post-test **(E, G)**.

**Figure S3.**
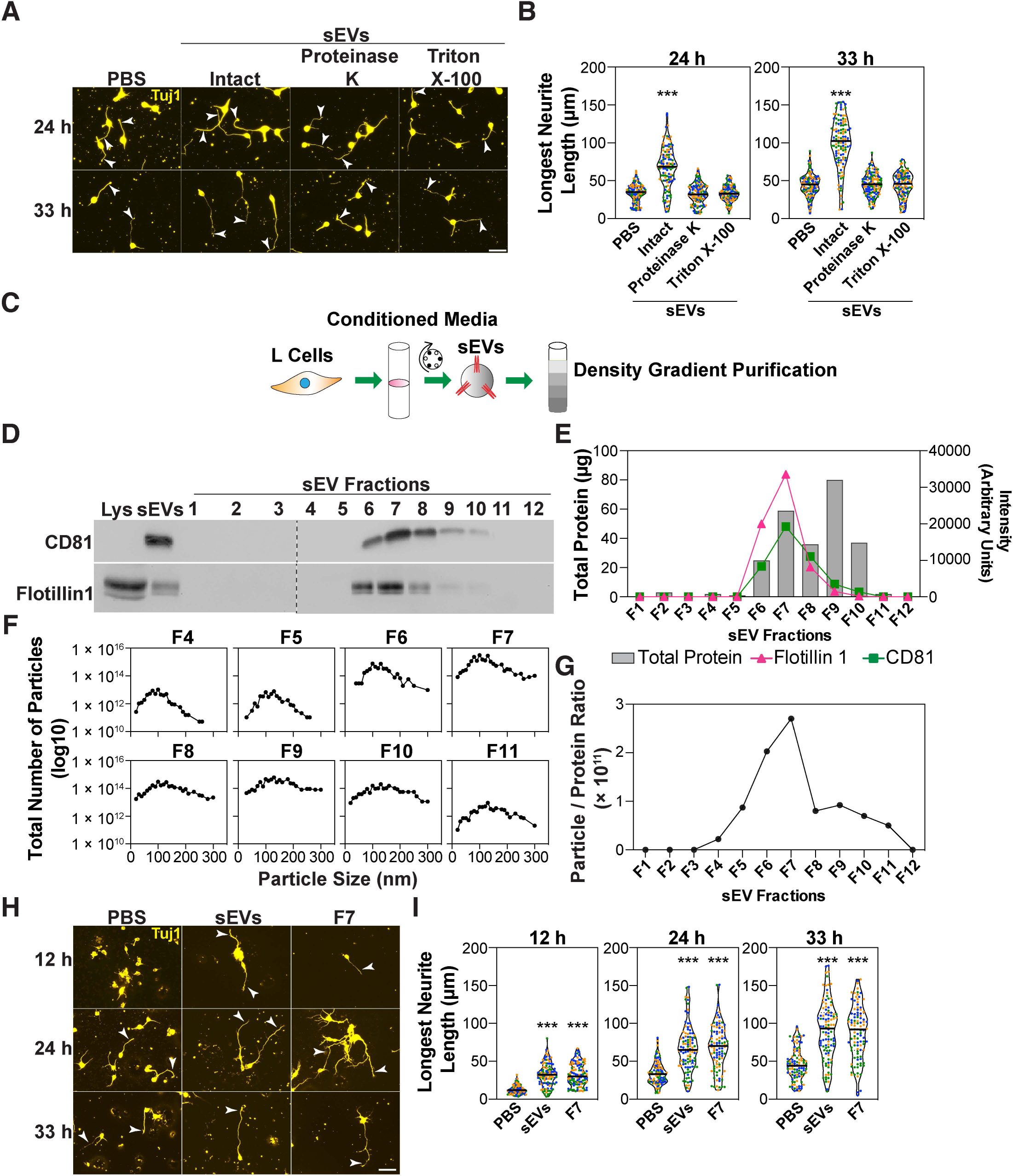
sEVs isolated using a discontinuous density gradient promote the growth of the longest neurite. Related to Fig. 1. **(A-B, H-I)** Dissociated E15.5-16.5 cortical neurons were treated with the intact sEVs, Proteinase K treated and Triton X-100 treated sEV pellet **(A-B)** or sEV pellet and the F7 fraction (both at 2 × 10^8^ particles/mL) **(H-I)**, 4 h after plating. Neuronal morphology was examined in Tuj1 stained neurons at 24 and 33 h **(A-B)** or 12, 24 and 33 h **(H-I)**. Representative images **(A, H)** and quantifications **(B, I)** are shown. Arrowheads mark the longest neurite. Scale bar, 40 μm. **(C)** A schematic of the experimental set up. sEV pellets (100,000 x g) purified from the conditioned media (CM) of L cells using differential centrifugation were subjected to iodixanol discontinuous density gradient purification. (**D)** Twelve fractions were collected starting from the top of the discontinuous gradient and were analyzed by immunoblotting for the indicated EV markers. (**E)** Intensity of the bands of EV markers from **(D)** is plotted against the protein concentration. (**F)** Particle size distribution was determined using NTA. (**G)** Fraction 7 (F7) has the highest particle number/protein ratio. Neurite lengths are quantified from a minimum of 90 neurons per condition from 3 independent experiments and plotted as a violin plot with values from each experiment distinctly colored and the median marked by a black line. Statistical significance: ***p<0.001 using one-way ANOVA with Dunnett’s post-test.

**Figure S4.**
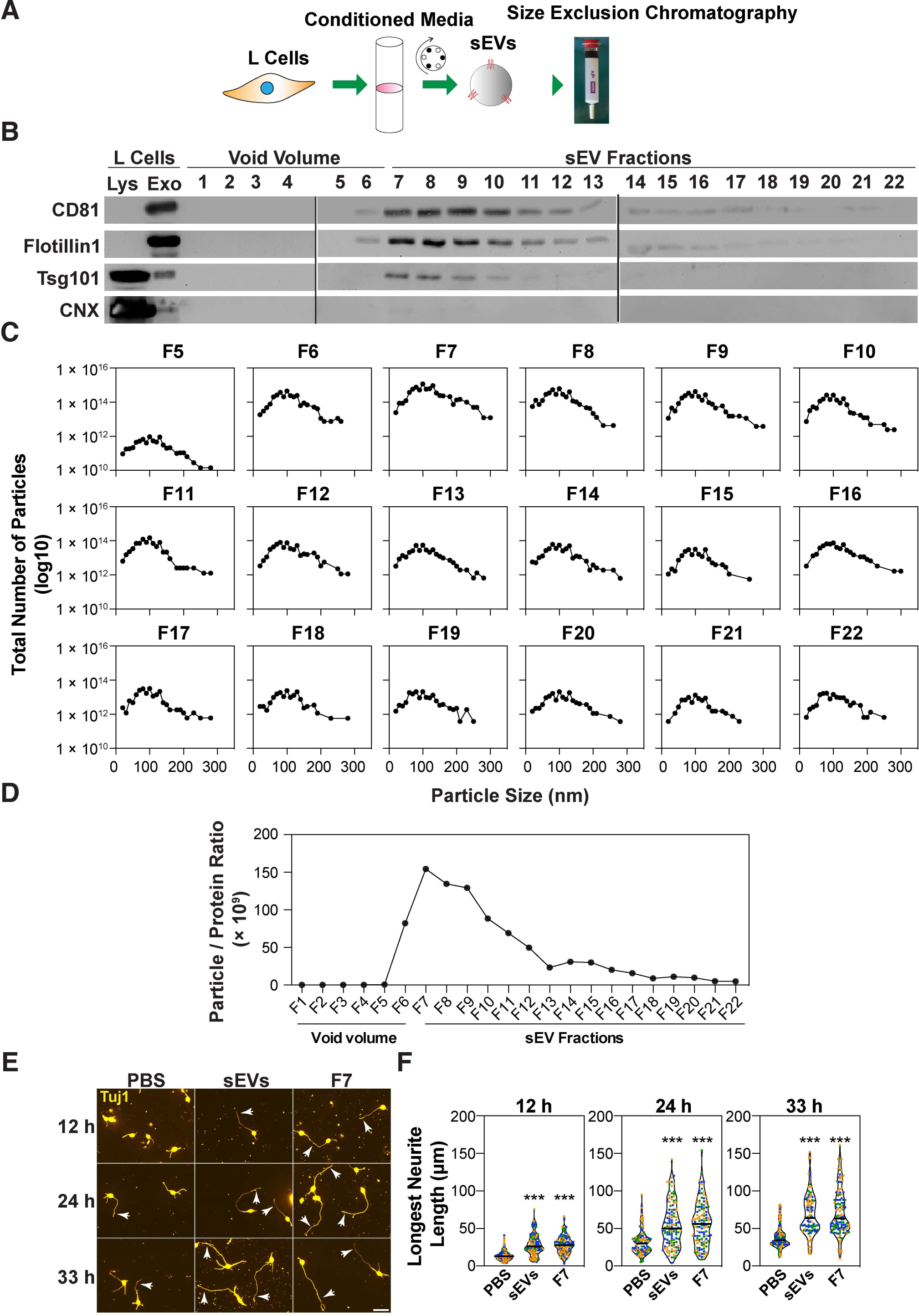
sEVs isolated using size exclusion chromatography promote the growth of the longest neurite. Related to Fig. 1. **(A)** A schematic illustration of experimental setup. sEVs purified from the CM of L cells were subjected to size exclusion chromatography. **(B)** A total of 22 fractions were collected and subjected to immunoblotting for EV markers. (**C)** Particle size was measured using NTA. **(D)** Fraction 7 (F7) has the highest particle number/protein ratio. **(E)** F7 fraction promotes the growth of the longest neurite. Dissociated E15.5-16.5 cortical neurons were treated with resuspended sEV pellet (5 μg/mL) and F7 fraction (5 μg/mL), 4 h after plating. Neuronal morphology was examined in Tuj1 stained neurons at various time points: 12, 24 and 33 h. Representative images are shown. Arrowheads mark the longest neurite. Scale bar, 40 μm. **(F)** The length of the longest neurite was quantified from a minimum of 90 neurons per condition from 3 independent experiments and plotted as a violin plot with values from each experiment distinctly colored and the median marked by a black line. Statistical significance: ***p<0.001 using one-way ANOVA with Dunnett’s post-test.

**Figure S5.**
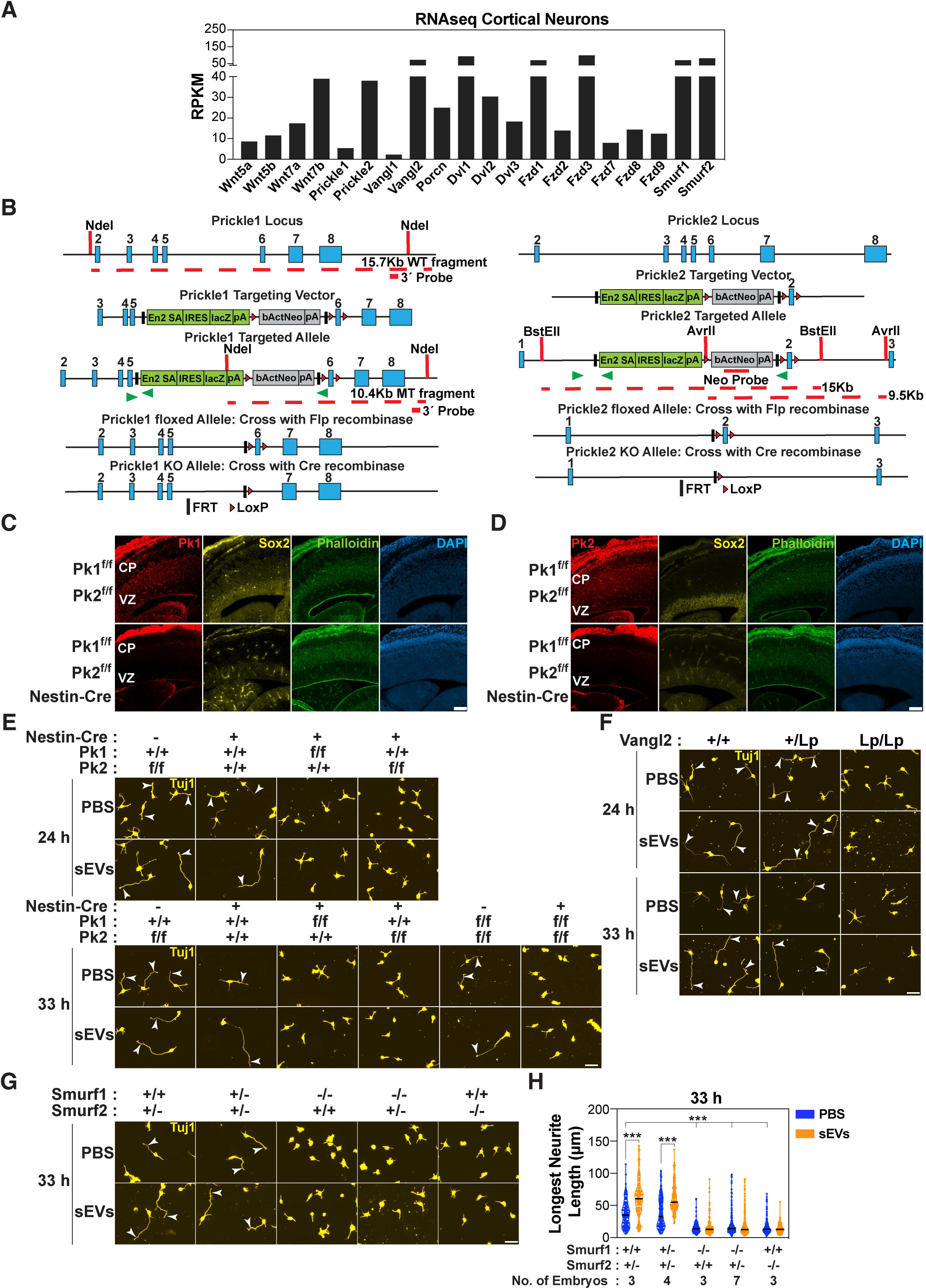
The PCP components, Pk1/2 and Vangl2, are required for sEV-induced growth of the longest neurite. Related to Fig. 2. **(A)** Expression of PCP genes and Wnts in cortical neurons. Total RNA was isolated from E15.5-16.5 cortical neurons after 33 h of *in vitro* culturing. An RNA sequence library was constructed, and high-quality reads were aligned to the *M. musculus* genome and expression of the indicated genes plotted. RPKM, Reads per kilobase of transcript per million mapped reads. (**B)** Schematic maps of the Pk1 and Pk2 wild type allele, targeting vector, targeted allele, floxed allele after crossing with Flp recombinase and knockout allele after crossing with Cre recombinase are shown. Green arrowheads indicate the position of PCR primers for genotyping. **(C, D)** Brain cryosections of E15.5 mouse embryo were stained with antibodies for Pk1 and Sox2 **(C)** or Pk2 and Sox2 **(D)** along with phalloidin. CP: cortical plate, VZ: ventricular zone. Scale bar, 100 μm. **(E-H)** Cortical neurons (E15.5-16.5) were isolated from Pk1 and Pk2 conditional knockout mice **(E)** or Vangl2 mutant littermates **(F)** or Smurf1 and Smurf2 knockout mice **(G, H)**, and treated with sEVs from L cells, 4 h after plating. Neurons were fixed at 24 and 33 h and stained with Tuj1. Representative images are shown. Arrowheads mark the longest neurite. Scale bar, 40 μm. **(H)** The length of the longest neurite was quantified from 30 neurons per embryo, and the total number of embryos analyzed was indicated below the genotypes. Statistical significance: ***p<0.001 using two-way ANOVA with Tukey’s post-test.

**Figure S6.**
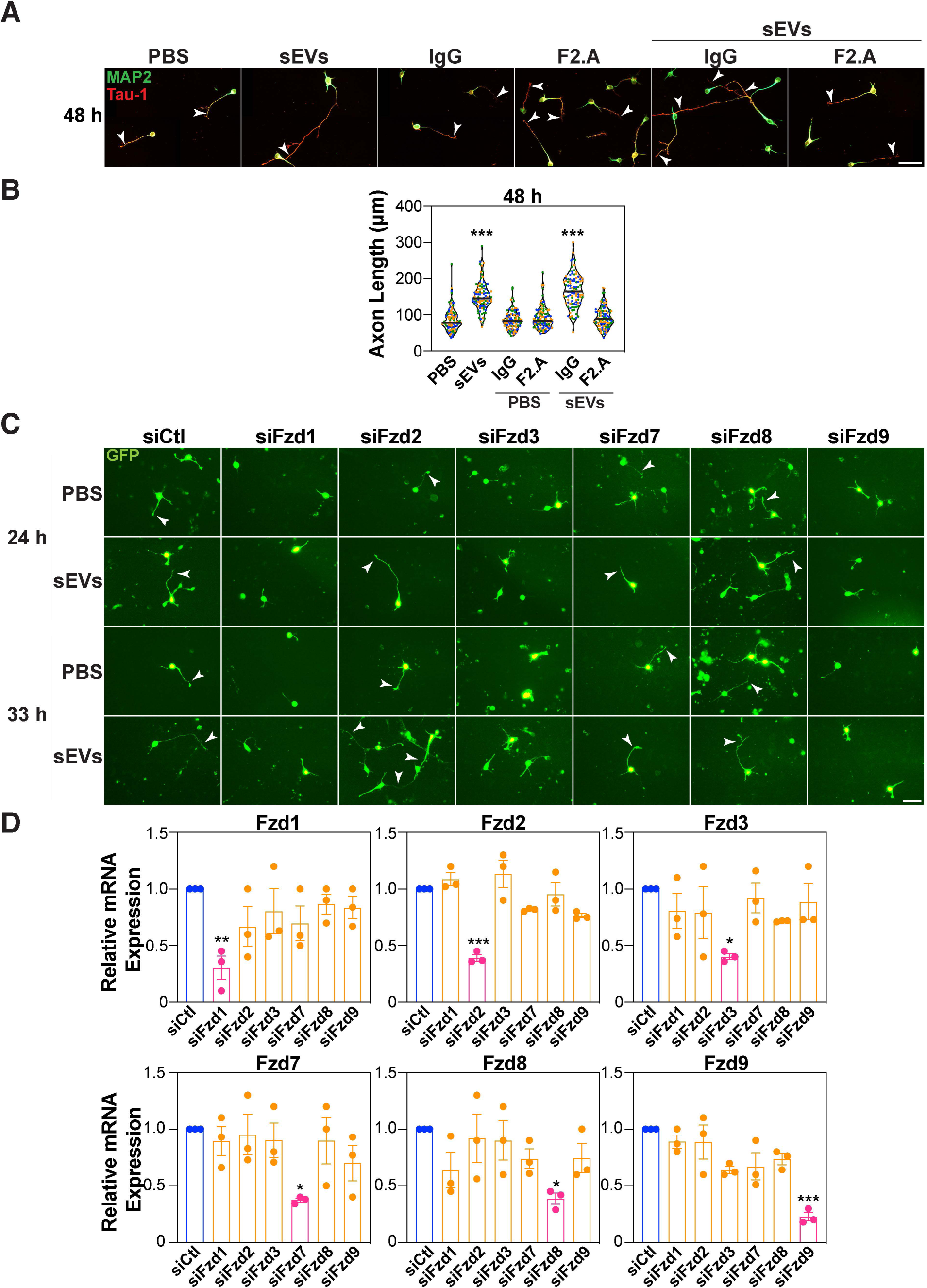
The PCP components, Fzds, are required for sEV-induced growth of the longest neurite. Related to Fig. 3. **(A-B)** Dissociated E15.5-16.5 mouse cortical neurons were treated with sEVs (5 μg/mL) from L cells and IgG or a Fzd blocking antibody, F2.A (100 nM), 24 h after plating. Neurons were fixed at 48 h, and neuronal morphology was examined in neurons immunostained for MAP2 (dendrites, green) and Tau-1 (axons, red). Representative images are shown **(A)**. Arrowheads mark Tau-1 positive axons. Scale bar, 50 μm. **(B)** The length of the Tau-1 positive axons was quantified from a minimum of 90 neurons per condition from 3 independent experiments and plotted as a violin plot with values from each experiment distinctly colored and the median marked by a black line. **(C)** Representative images of dissociated E15.5-16.5 cortical neurons electroporated with siRNA against Fzds (siFzds) or siControl (siCtl) along with a GFP-expressing plasmid and then treated with sEVs (5 μg/mL) from L cells, 4 h after plating. Arrowheads mark the longest neurite. Scale bar, 40 μm. **(D)** Knockdown efficiency for Fzds was determined in GFP-positive neurons isolated by FACS. Relative mRNA expression was determined by qPCR and is plotted as the mean ± SEM from 3 independent experiments. Statistical significance: *p<0.05, **p<0.01, ***p<0.001 using one-way ANOVA with Dunnett’s post-test.

**Figure S7.**
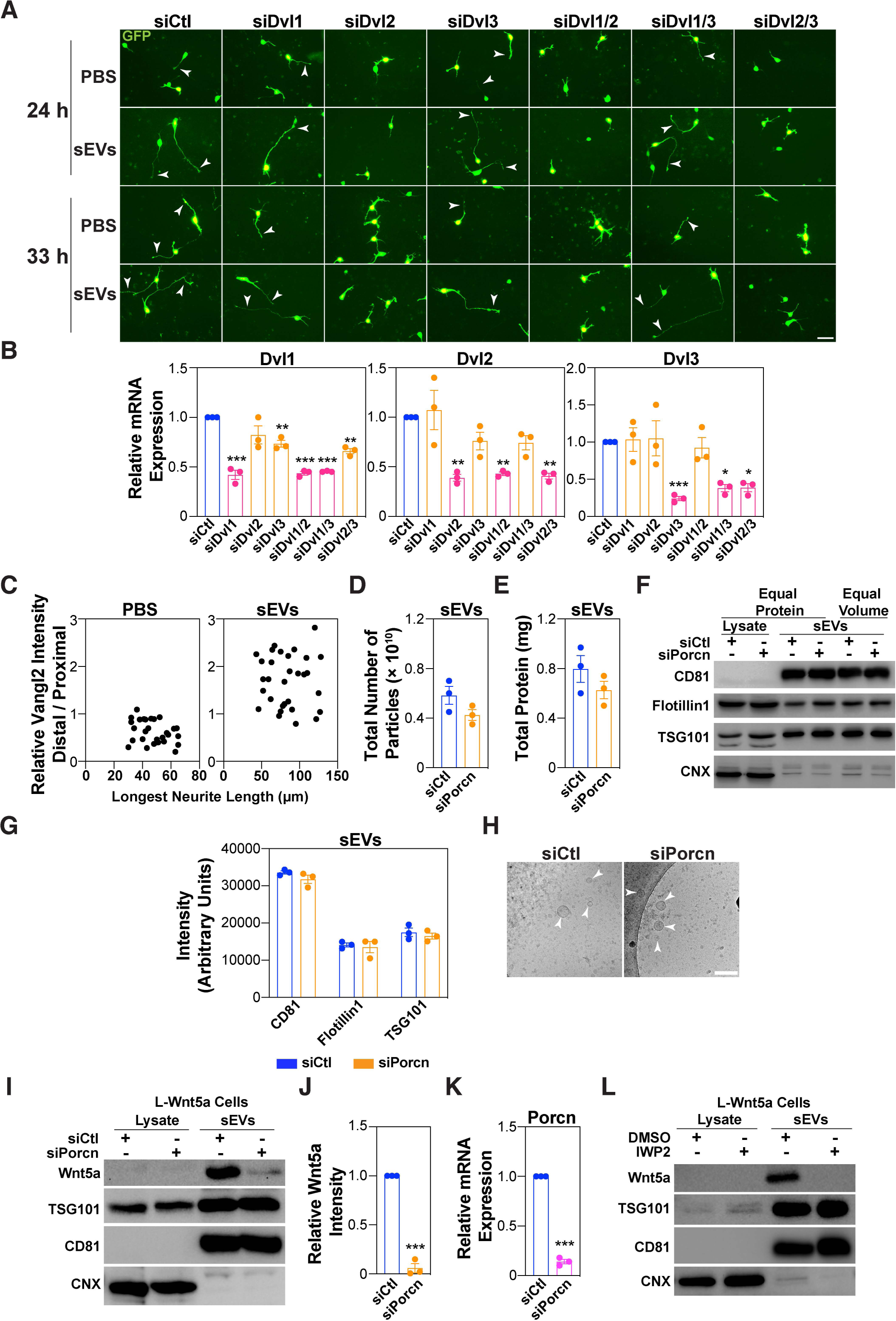
Related to Fig. 3 and 4. **(A)** Representative images of dissociated E15.5-16.5 cortical neurons electroporated with siRNA against Dvls (siDvls) or siControl (siCtl) along with a GFP-expressing plasmid and then treated with sEVs (5 μg/mL) from L cells, 4 h after plating. Arrowheads mark the longest neurite. Scale bar, 40 μm. **(B)** Knockdown efficiency for Dvls was determined in GFP-positive neurons isolated by FACS. Relative mRNA expression was determined by qPCR and is plotted as the mean ± SEM from 3 independent experiments. (**C)** The ratio of distal/proximal Vangl2 intensity as a function of the longest neurite length in neurons treated with PBS or sEVs is quantified from 30 neurons from 3 independent experiments. (**D)** The total number of particles in the sEV pellet (100,000 x g) were measured using NTA. (**E)** The total amount of protein in the sEV pellet was measured by Bradford assay. (**F)** Characterization of sEVs derived from porcupine-deficient L cells. Representative immunoblot of the cell lysate and 100,000 x g pellet (sEVs) analyzed for EV markers and calnexin (CNX). (**G)** Quantification of **(F)**. sEVs were collected from an equal number of cells, and equal volumes of sEV pellets (right two lanes) were used for comparison. Protein bands were quantified using ImageJ. **(H)** Representative TEM images of sEV pellet. Arrowheads indicate round vesicles. Scale bar, 200 nm. **(I-L)** Porcupine inhibition blocks Wnt5a secretion in sEVs. L-Wnt5a cells were treated with siCtl or siPorcupine (siPorcn) (**I-K**) and DMSO or IWP2 (10 μM) **(L)** followed by isolation of sEVs. A representative immunoblot of sEVs and lysates indicating the level of Wnt5a, EV markers and CNX is shown **(I, L)**. **(J)** Quantification of **(I)**. Protein bands were quantified using ImageJ. **(K)** Knockdown efficiency of Porcupine in L-Wnt5a cells. Values are plotted as the mean ± SEM from 3 independent experiments **(B, D, E, G, K, L)**. Statistical significance: *p<0.05, **p<0.01, ***p<0.001 using unpaired t-test **(D, E, K, L)**, one-way ANOVA with Dunnett’s post-test **(B)**, or two-way ANOVA with Tukey’s post-test **(G)**.

**Figure S8.**
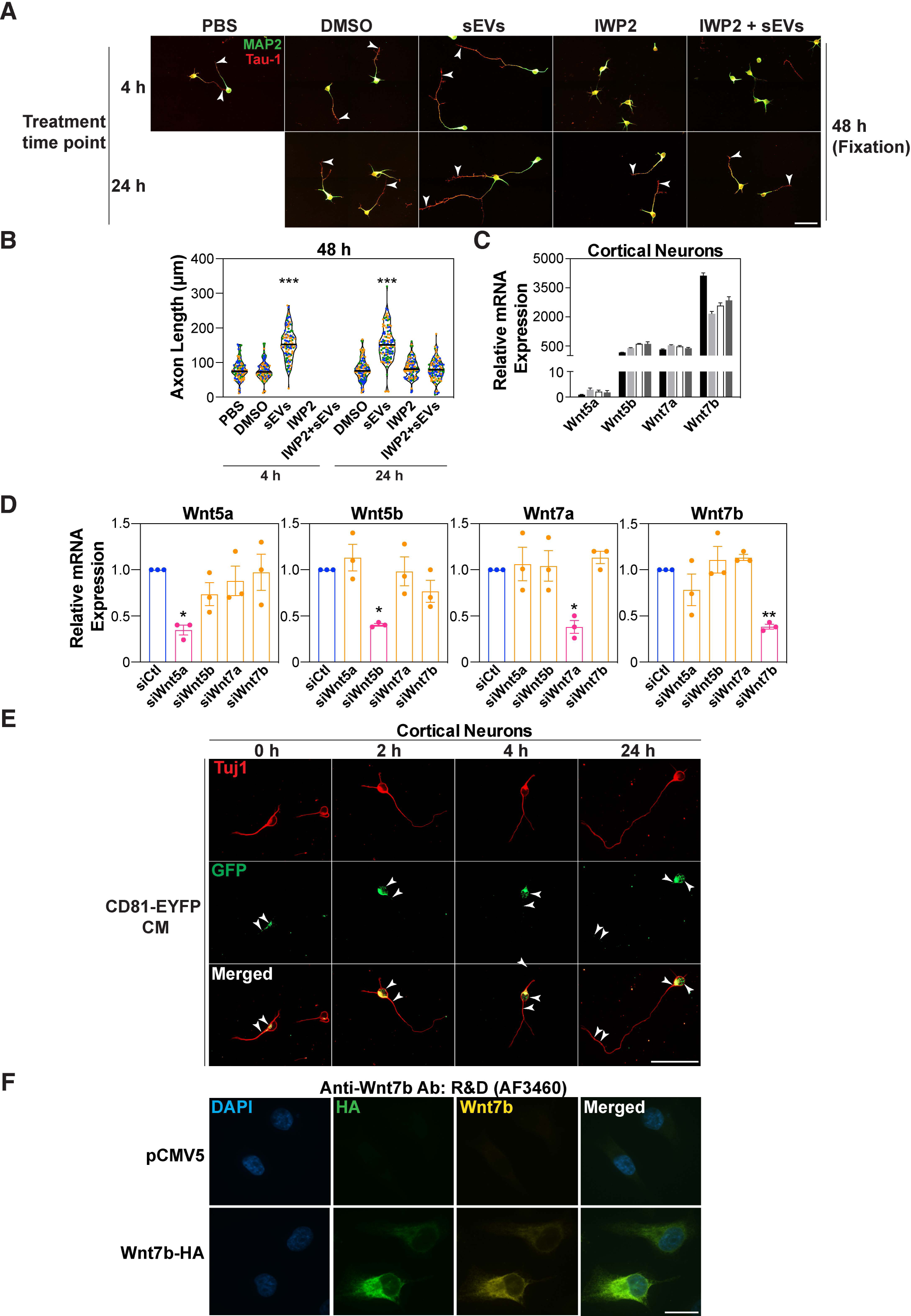
Related to Fig. 4 and 5. **(A-B)** Dissociated E15.5-16.5 mouse cortical neurons were treated with sEVs (5 μg/mL) from L cells and DMSO or IWP2 (10 μM), 4 h or 24 h after plating. Neurons were fixed at 48 h, and neuronal morphology was examined in neurons immunostained for MAP2 (dendrites, green) and Tau-1 (axons, red). Representative images are shown **(A)**. Arrowheads mark Tau-1 positive axons. Scale bar, 50 μm. **(B)** The length of the Tau-1 positive axons was quantified from a minimum of 90 neurons per condition from 3 independent experiments and plotted as a violin plot with values from each experiment distinctly colored and the median marked by a black line. For IWP2-treated neurons at 4 h, a comparable number of data points were not available due to lack of Tau-1 positive axons. **(C)** Expression of Wnts in cortical neurons. RNA was isolated from cortical neurons at 0, 12, 24 and 33 h of culturing. A representative plot indicating mRNA expression relative to Wnt5a, determined by qPCR from 2 independent experiments is shown. **(D)** Efficiency of siRNA-mediated knockdown of Wnts. RNA was isolated from GFP-positive neurons after FACS. Relative mRNA expression was determined by qPCR. **(E)** sEVs can be internalized by neurons. Cortical neurons were treated with 10X concentrated conditioned media (CM) from L cells stably expressing CD81-EYFP, 24 h after plating for 30 min. Neurons were washed and subsequently treated with regular complete media for 0, 2, 4 and 24 h. Representative images of neurons immunostained with GFP and Tuj1 from 30 neurons from 3 independent experiments are shown. Arrowheads indicate the localization of sEVs. Scale bar, 40 μm **(F)** Wnt7b antibody characterization. MDA-MB-231 cells were transfected with 2 μg of either pCMV5 control or C-terminal HA-tagged Wnt7b (Wnt7b-HA). Cells were fixed after 48 h and immunostained with HA (green) and Wnt7b (yellow) antibodies. Scale bar, 20 μm. Statistical Significance: *p<0.05, **p<0.01, ***p<0.001 using one-way ANOVA with Dunnett’s post-test **(B, D)**.

**Figure S9.**
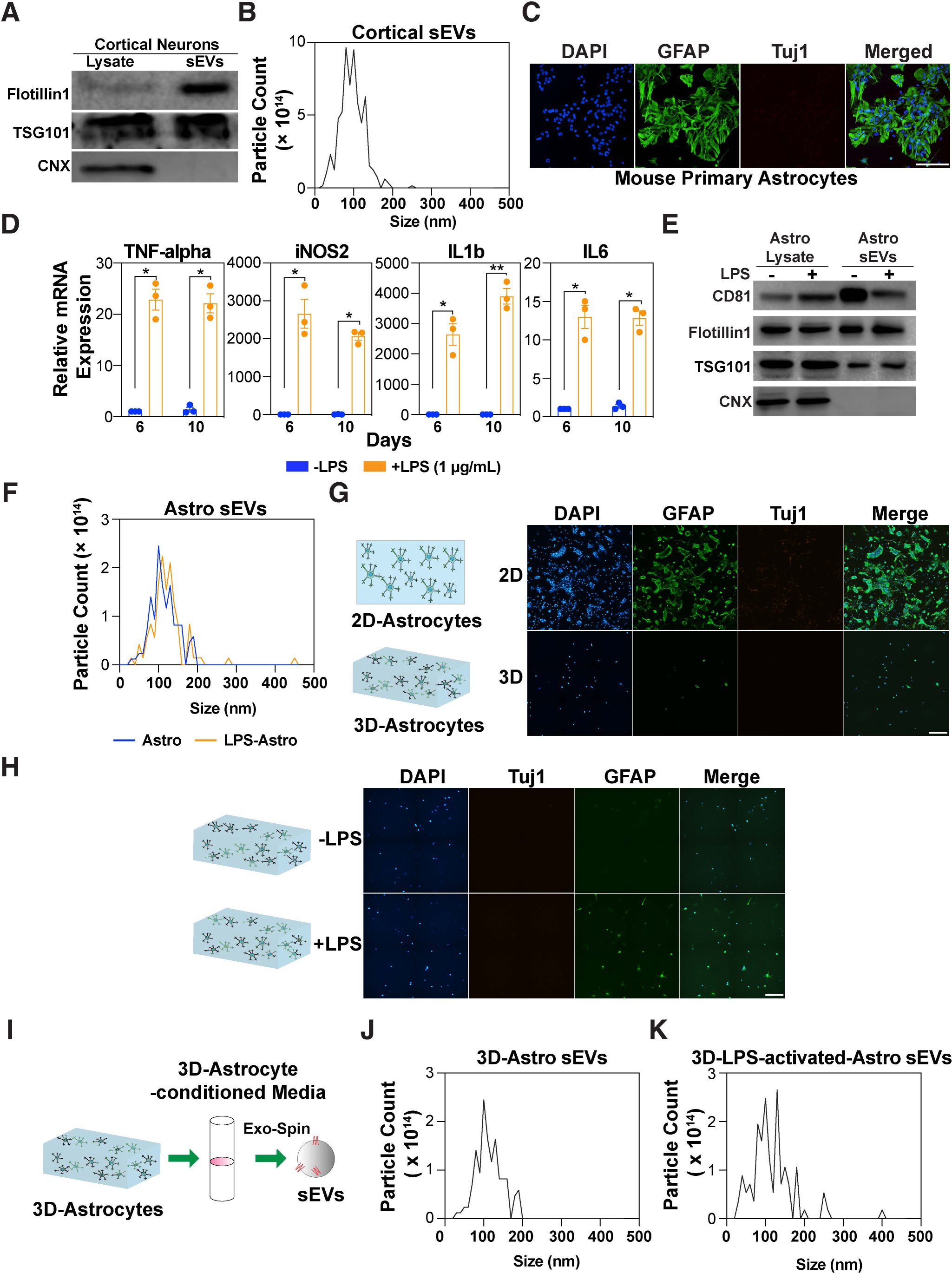
Related to Fig. 6. **(A)** Immunoblot of sEVs and lysates from cortical neurons. **(B)** NTA of the sEV pellet. **(C)** Purity of primary astrocytes cultures. Enriched cultures of astrocytes from P1-P4 mouse pups were immunostained for the astrocyte marker, GFAP, and the neuronal marker Tuj1. Scale bar, 200 μm. **(D)** LPS activates mouse primary astrocytes. Astrocytes were treated with LPS (1 μg/mL), and RNA was isolated after 6 or 10 days. Relative expression of pro-inflammatory genes was measured by qPCR. **(E)** Immunoblot of lysates and the sEV pellet from primary astrocytes. **(F)** NTA of the sEV pellet. **(G)** Astrocytes are activated upon culturing in 2D plates. Representative immunofluorescence images of astrocytes grown either on a glass surface or in a soft collagen gel. Scale bar, 200 μm. **(H)** LPS activates astrocytes grown in 3D-collagen gels. Astrocytes were treated with LPS (1 μg/mL), fixed after 48 h and subsequently immunostained with GFAP and Tuj1 antibodies. Representative images are shown. Scale bar, 200 μm. **(I)** A schematic representation of sEV purification from astrocytes grown in a soft 3D-collagen gel. CM was collected after 6 days and subsequently sEVs were purified using Exo-spin^TM^ columns. **(J and K)** NTA of the sEVs. NTA plots are from 1 representative experiment out of 3 independent purifications **(B, F, J, K)**. Values are plotted as the mean ± SEM from 3 independent experiments **(D)**. Statistical significance: *p<0.05, **p<0.01 using two-way ANOVA with Tukey’s post-test **(D)**.

**Table S1:**
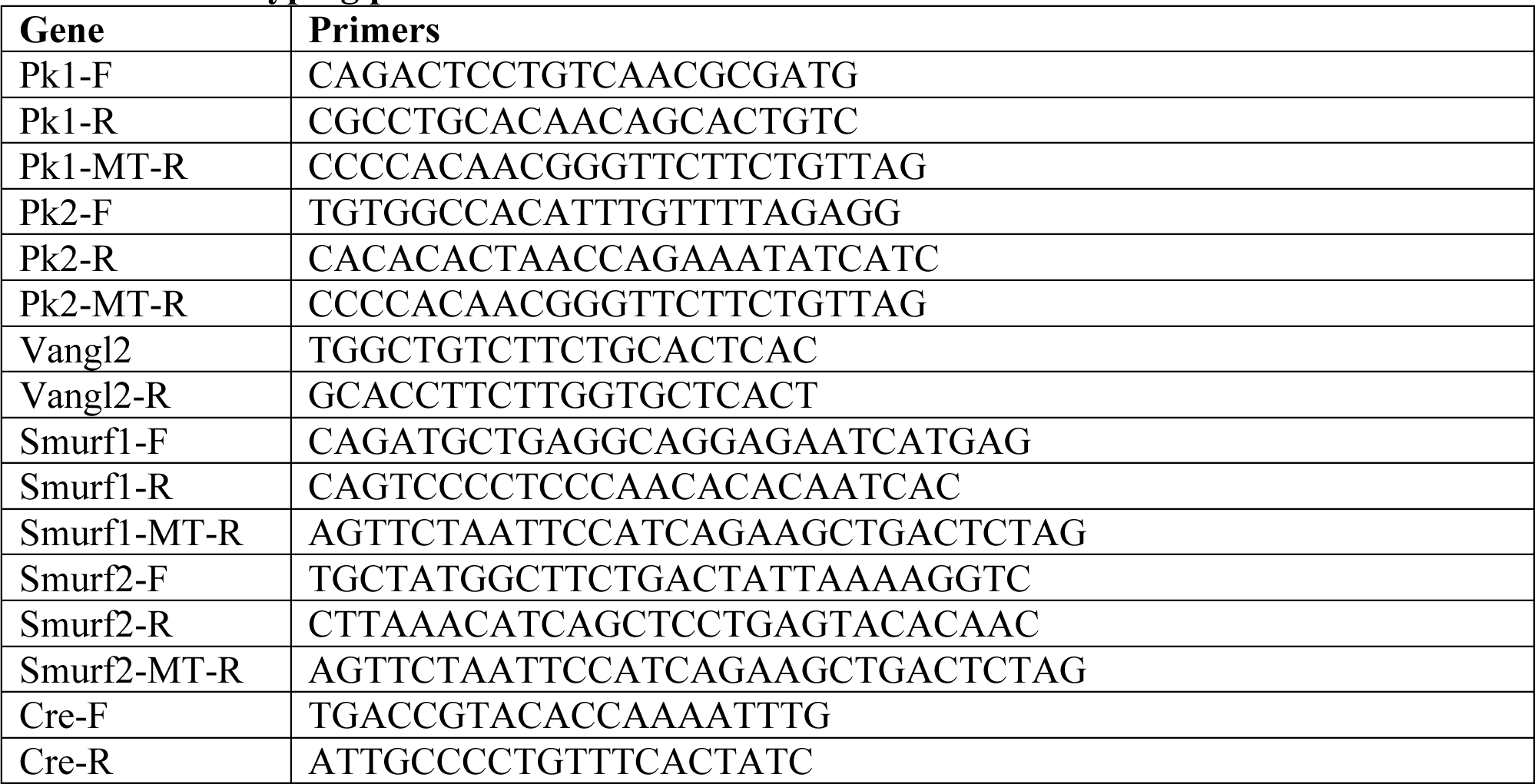
Genotyping primers.

**Table S2:**
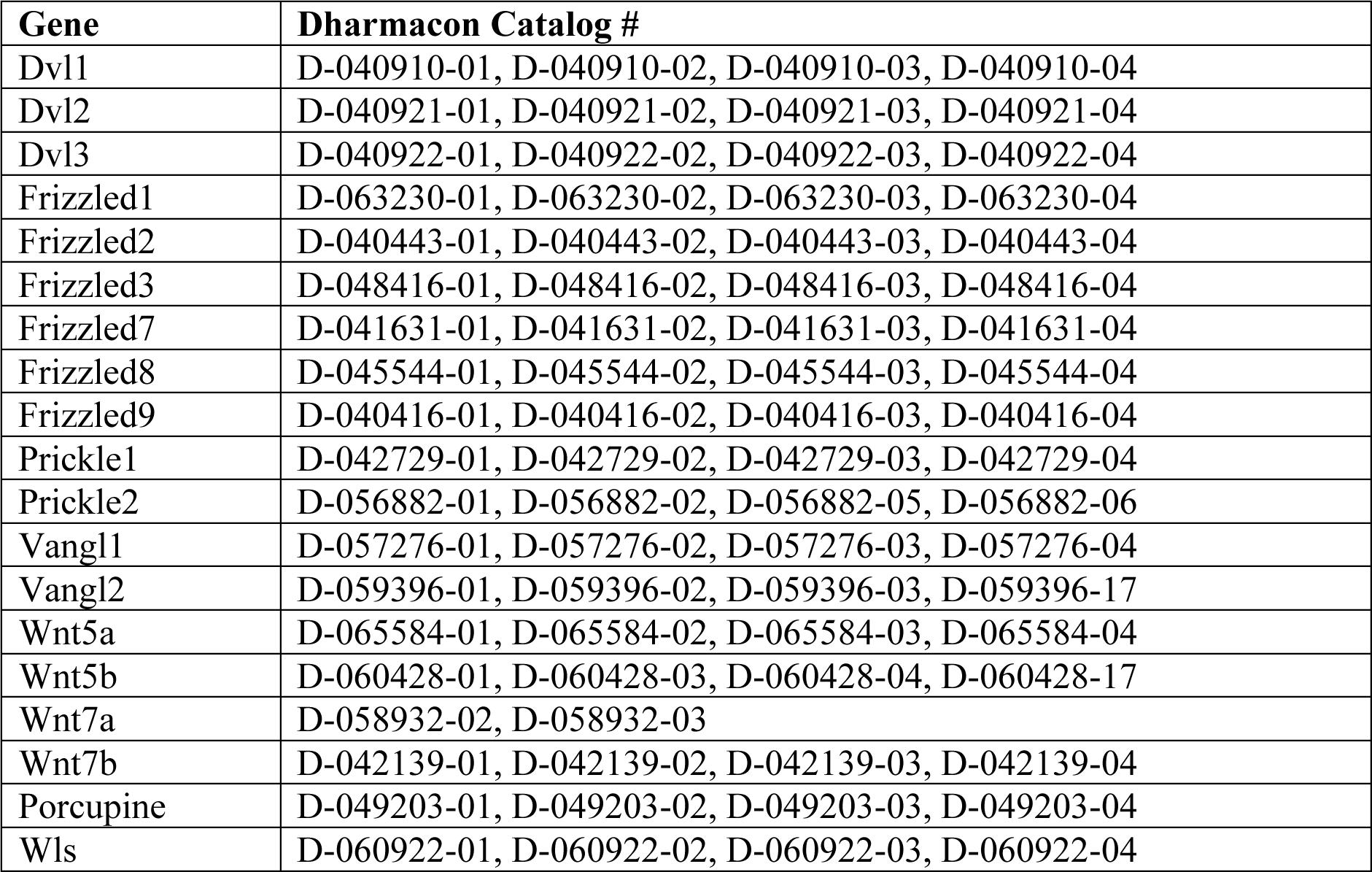
List of siRNAs.

| Gene | Dharmacon Catalog # |
| --- | --- |
| Dvl1 | D-040910-01, D-040910-02, D-040910-03, D-040910-04 |
| Dvl2 | D-040921-01, D-040921-02, D-040921-03, D-040921-04 |
| Dvl3 | D-040922-01, D-040922-02, D-040922-03, D-040922-04 |
| Frizzled1 | D-063230-01, D-063230-02, D-063230-03, D-063230-04 |
| Frizzled2 | D-040443-01, D-040443-02, D-040443-03, D-040443-04 |
| Frizzled3 | D-048416-01, D-048416-02, D-048416-03, D-048416-04 |
| Frizzled7 | D-041631-01, D-041631-02, D-041631-03, D-041631-04 |
| Frizzled8 | D-045544-01, D-045544-02, D-045544-03, D-045544-04 |
| Frizzled9 | D-040416-01, D-040416-02, D-040416-03, D-040416-04 |
| Prickle1 | D-042729-01, D-042729-02, D-042729-03, D-042729-04 |
| Prickle2 | D-056882-01, D-056882-02, D-056882-05, D-056882-06 |
| Vangl1 | D-057276-01, D-057276-02, D-057276-03, D-057276-04 |
| Vangl2 | D-059396-01, D-059396-02, D-059396-03, D-059396-17 |
| Wnt5a | D-065584-01, D-065584-02, D-065584-03, D-065584-04 |
| Wnt5b | D-060428-01, D-060428-03, D-060428-04, D-060428-17 |
| Wnt7a | D-058932-02, D-058932-03 |
| Wnt7b | D-042139-01, D-042139-02, D-042139-03, D-042139-04 |
| Porcupine | D-049203-01, D-049203-02, D-049203-03, D-049203-04 |
| Wls | D-060922-01, D-060922-02, D-060922-03, D-060922-04 |

**Table S3:** List of antibodies.

| Reagent or Resource | Source | Identifier |
| --- | --- | --- |
| **Antibodies** |  |  |
| Mouse anti-MAP-2 (Clone AP20, 1:500) | Millipore | MAB3418 |
| Mouse anti-Tubulin $\beta$ 3 (Clone Tuj1, 1:3000) | Biolegend | 801202 |
| Mouse anti-Tau-1 (Clone PC1C6, 1:300) | Millipore | MAB3420 |
| Mouse anti-TSG101 (1:1500) | GeneTex | GTX70255 |
| Mouse anti-Flotillin-1 (1:1500) | BD Transduction | 618021 |
| Mouse anti-CD81 (B-11, 1:1500) | Santa Cruz | sc-166029 |
| Mouse anti-Prickle1 (F-5, 1:100) | Santa Cruz | Sc-393034 |
| Mouse anti-Prickle2 (Clone 3B4.1, 1:100) | Millipore | MABN1529 |
| Rabbit anti-GFAP (1:500) | Dako | Z0334 |
| Rabbit anti-GFP (1:400) | Invitrogen | A11122 |
| Rabbit anti-β III Tubulin (1:500) | Abcam | ab18207 |
| Rabbit anti-Calnexin (1:2000) | (Jackson *et al.*, 1994)<sup>1</sup> | N/A |
| Rat anti-Vangl2 (Clone 2G4, 1:200) | Millipore | MABN750 |
| Rat anti-HA (1:1000) | Roche | 11867423001 |
| Goat anti-Wnt7b (1:200) | R&D Systems | AF3460 |
| Goat anti-Wnt5a (1:1000) | R&D Systems | AF645 |
| Goat anti-GFP (1:1000) | Rockland | 600-101-215 |
| Donkey anti-goat DyLight<sup>TM</sup> 488 (1:1000) | Thermo Fisher Scientific | SA5-10086 |
| Donkey anti-goat Alexa Fluor<sup>TM</sup> 647 (1:1000) | Thermo Fisher Scientific | A21447 |
| Goat anti-Mouse Alexa Fluor<sup>®</sup> 488 (1:1000) | Invitrogen | A11029 |
| Goat anti-Rabbit Alexa Fluor<sup>®</sup> 488 (1:1000) | Invitrogen | A11034 |
| Goat anti-Mouse Alexa Fluor<sup>TM</sup> 488 (1:500) | Thermo Fisher Scientific | A21121 |
| Goat anti-Rat Alexa Fluor<sup>TM</sup> 488 (1:1000) | Thermo Fisher Scientific | A11006 |
| Goat anti-Mouse Alexa Fluor<sup>TM</sup> 568 (1:500) | Thermo Fisher Scientific | A21134 |
| Goat anti-Mouse Alexa Fluor<sup>®</sup> 594 (1:1000) | Invitrogen | A11032 |
| Goat anti-Rabbit Alexa Fluor<sup>®</sup> 594 (1:1000) | Invitrogen | A11012 |
| Goat anti-rabbit Alexa Fluor<sup>TM</sup> 647 (1:1000) | Thermo Fisher Scientific | A21244 |
| Donkey anti-Goat IgG-HRP (1:5000) | Jackson ImmunoResearch | 705-035-147 |
| Donkey anti-Mouse IgG-HRP (1:5000) | Jackson ImmunoResearch | 715-035-150 |
| Donkey anti-Rabbit IgG-HRP (1:5000) | Jackson ImmunoResearch | 711-035-152 |
| IgG F2.A | (Pavlovic *et al.*, 2018)<sup>2</sup> | N/A |
| IgG | (Enderle *et al.*, 2021)<sup>3</sup> | N/A |

<sup>1</sup>Jackson MR, Cohen-Doyle MF, Peterson PA, Williams DB (1994) Regulation of MHC class I transport by the molecular chaperone, calnexin (p88, IP90). *Science* 263: 384-387.
<sup>2</sup>Pavlovic Z, Adams JJ, Blazer LL, Gakhal AK, Jarvik N, Steinhart Z, Robitaille M, Mascall K, Pan J, Angers S *et al* (2018) A synthetic anti-Frizzled antibody engineered for broadened specificity exhibits enhanced anti-tumor properties. *MAbs* 10: 1157-1167.
<sup>3</sup>Enderle L, Shalaby KH, Gorelik M, Weiss A, Blazer LL, Paduch M, Cardarelli L, Kossiakoff A, Adams JJ, Sidhu SS (2021) A T cell redirection platform for co-targeting dual antigens on solid tumors. *MAbs* 13: 1933690.

**Table S4:**
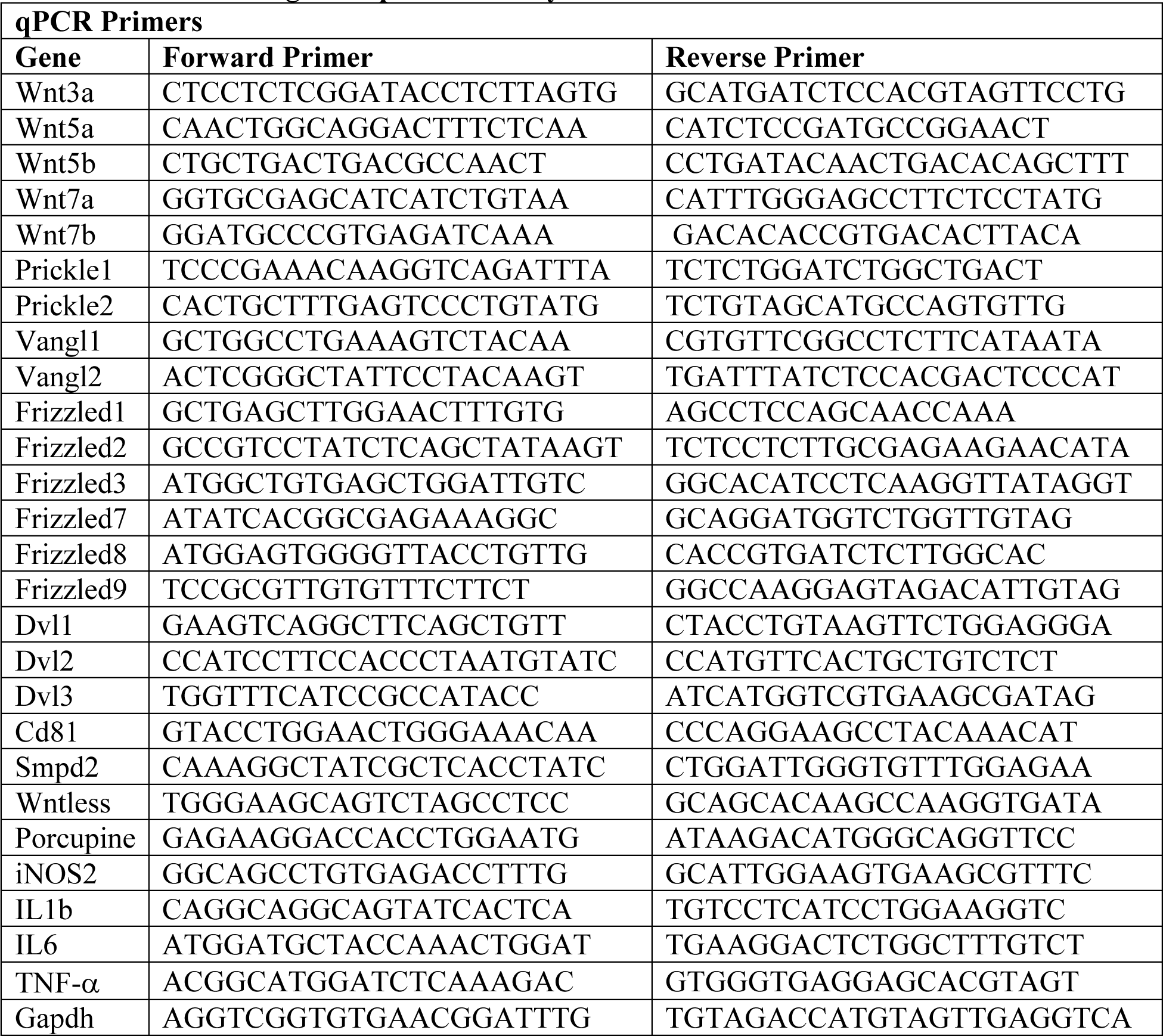
Primers for gene expression analysis.

